# Anatomically constrained and curated cerebellar tractography (ACCURAT): an open framework and a pathway-specific neuroanatomical reference

**DOI:** 10.64898/2026.05.02.722413

**Authors:** Jon Haitz Legarreta, Richard J. Rushmore, Edward H. Yeterian, Nikos Makris, Yogesh Rathi, Lauren J. O’Donnell

## Abstract

Cerebellar pathways form extensive structural circuits linking the cerebellum with the brainstem, thalamus, and cerebrum, underlying motor, cognitive, and affective functions. Diffusion MRI tractography provides the only non-invasive method for mapping these pathways in vivo, but reconstruction of cerebellar connectivity remains challenging due to crossing fibers, peduncular bottlenecks, decussations, multi-synaptic circuits, and numerous small nuclei that define pathway origins and terminations. Here we introduce <u>A</u>natomically <u>C</u>onstrained and <u>CURA</u>ted <u>T</u>ractography (ACCURAT), an open framework for reconstructing cerebellar pathways from diffusion MRI using anatomical priors and rule-based streamline queries. ACCURAT combines anatomical segmentation, densely seeded tractography, and vertex-level evaluation of anatomical constraints along streamline trajectories, enabling the isolation of pathway segments within specific nuclei while preventing their propagation across synaptic boundaries. To define these constraints, we provide a concise, pathway-by-pathway synthesis of cerebellar connectional anatomy based on experimental tract-tracing literature and organized for tractography applications. We identify pathway-specific origins, trajectories, terminations, decussation patterns, and tractography challenges, and use this information to inform tractography-ready cerebellar pathway definitions. Using ultra-high-resolution submillimeter diffusion MRI (0.76 mm gSlider acquisition) from healthy participants, we reconstruct multiple extrinsic and intrinsic cerebellar pathways, including specific components of the inferior, middle, and superior cerebellar peduncles; challenging decussating pathways such as the olivocerebellar and dentato-olivary projections; and intrinsic cerebellar pathways, including Purkinje corticonuclear projections and intracortical parallel fibers. ACCURAT generalizes across tractography algorithms, producing comparable reconstructions with both probabilistic parallel transport tractography and deterministic unscented Kalman filter tractography. Together, the ACCURAT framework and accompanying neuroanatomical reference provide an anatomically grounded, tractography-oriented resource for reconstructing cerebellar pathways in vivo and for supporting future development and evaluation of cerebellar tractography methods.

## 1. Introduction

The cerebellum plays a critical role in motor coordination, cognition, language, emotion, and perception (Buckner, 2013; Strick et al., 2009; B. Wang et al., 2025). Consistent with these diverse functions, the cerebellum participates in extensive anatomical circuits linking the cerebellar cortex with the brainstem, spinal cord, and thalamus, as well as indirectly with the cerebral cortex. These circuits form a complex structural connectome that supports sensorimotor integration, motor learning, and higher cognitive functions. Disruptions of cerebellar connectivity have been implicated in numerous neurological and psychiatric disorders, including autism, schizophrenia, Parkinson’s disease, and Alzheimer’s disease (Fatemi et al., 2012; Gellersen et al., 2021; Jacobs et al., 2018; Phillips et al., 2015; Reeber et al., 2013; Sathyanesan et al., 2019; Wu & Hallett, 2013).

The structural organization of the cerebellum includes both extrinsic pathways, which connect the cerebellum with the brainstem and cerebrum, and intrinsic pathways, which comprise the intracerebellar circuitry. The principal extrinsic pathways travel through the three cerebellar peduncles. The inferior cerebellar peduncle (ICP) primarily carries afferent input from the spinal cord and inferior olive to the cerebellar cortex. The middle cerebellar peduncle (MCP) conveys afferent (input) projections from the pontine nuclei that relay cortical information to the cerebellum. The superior cerebellar peduncle (SCP) is the major efferent (output) pathway of the cerebellum and projects from the deep cerebellar nuclei to the red nucleus and thalamus. Within the cerebellum itself, intrinsic connections include Purkinje cell axons, which constitute the primary output of the cerebellar cortex to the deep cerebellar nuclei, and parallel fibers, the axons of granule cells that run within the cerebellar cortex along the cerebellar folia and provide a major excitatory input to Purkinje cells.

Diffusion MRI (dMRI) tractography provides the only non-invasive method for mapping these cerebellar connections in vivo, as described in several recent reviews (Beez et al., 2021; Habas & Manto, 2018; Harrison et al., 2021; Lundell & Steele, 2024). In the cerebrum, substantial progress has been made in improving tractography accuracy through anatomical constraints (Schilling et al., 2020; R. E. Smith et al., 2012), outlier removal (F. Zhang et al., 2018; Z. Zhang et al., 2018), rule-based bundle identification (Wassermann et al., 2016), and machine learning bundle segmentation (Barati Shoorche et al., 2025). For example, anatomically constrained tractography (ACT) uses cortex, white matter, and subcortical gray matter segmentations to guide streamline propagation and termination, typically ensuring that streamlines propagate through white matter and terminate upon reaching gray matter structures (R. E. Smith et al., 2012). The ACT constraints operate primarily at the level of tissue classes rather than pathway-specific anatomical rules. Another related approach, White Matter Query Language (WMQL), provides a rule-based framework for extracting white matter pathways using logical queries over anatomical regions of interest (ROIs), but it operates at the level of entire streamlines and cannot apply constraints to individual streamline vertices (Wassermann et al., 2016). These approaches incorporate anatomical knowledge of pathway origins, terminations, and trajectories to reduce the well-known prevalence of tractography false positives and false negatives (Maier-Hein et al., 2017; Schilling et al., 2019), which commonly arise from challenges such as crossing fibers (Jeurissen et al., 2013), bottlenecks (Schilling et al., 2022), and other ambiguities. However, existing approaches focus on cerebral pathways and do not address the anatomical organization of cerebellar circuits or their associated nuclei. These strategies have not yet been systematically extended to the cerebellar connectome.

Mapping cerebellar connectivity with tractography presents several unique challenges. First, there are challenges in mapping crossing axonal pathways within cerebellar white matter (Takahashi et al., 2013) and bottlenecks due to densely packed peduncles (Harrison et al., 2021). Second, many cerebellar pathways involve decussations or multi-synaptic circuits (Habas & Manto, 2018; Kwon et al., 2011), which complicate the identification of pathway endpoints and the segmentation of bundles. Third, cerebellar and brainstem pathways involve numerous small nuclei that serve as origins or terminations of axonal projections, making it difficult to define and enforce anatomically correct start and termination regions, especially at typical dMRI resolutions (Lundell & Steele, 2024). Because diffusion MRI does not encode sites of pathway termination (synapses) (Aydogan et al., 2018; C.-H. Yeh et al., 2020), tractography algorithms may propagate streamlines through nuclei where some pathways terminate. As a result, streamlines may continue along downstream pathways and effectively trace multi-synaptic circuits as single trajectories. Accurate reconstruction of cerebellar connectivity therefore requires anatomical priors specifying where streamlines should begin or terminate, and where they should be allowed to pass.

Several specific examples illustrate these difficulties. For example, the MCP is frequently reconstructed as a direct connection between cerebellar hemispheres (Carrasco-Guerrero et al., 2025; Leitner et al., 2015; Warrington et al., 2022; Yin et al., 2023; F. Zhang et al., 2018) rather than as two separate projections originating in the left and right pontine nuclei and connecting to the opposite cerebellar hemisphere (Martinez et al., 2026). Similarly, tractography of the SCP often extends into cerebral gray matter (Habas & Manto, 2018), effectively tracing cerebello-thalamo-cortical pathways across thalamic synapses rather than isolating the SCP itself. The SCP has synapses in the red nucleus and thalamus, although some axons traverse the red nucleus en route to the thalamus. Tractography of the ICP is almost universally presented as ipsilateral (Habas & Manto, 2018; van Baarsen et al., 2016). However, a major component of the ICP, the olivocerebellar pathway, originates in the contralateral inferior olivary nucleus and decussates before entering the cerebellum; reconstruction of this pathway remains challenging (Yin et al., 2023). Intrinsic cerebellar pathways pose additional challenges. Tractography studies of Purkinje fibers are often restricted to connections with the dentate nucleus (Jeong et al., 2014), overlooking cerebellar cortical projections to the other deep cerebellar nuclei. Parallel fibers—the most numerous axons in the cerebellum—are also frequently ignored because they lie within the cerebellar cortex, whereas tractography is typically restricted to white matter.

To address these challenges, we introduce **Anatomically Constrained and CURAted Tractography (ACCURAT)**, a framework that integrates anatomical priors with rule-based streamline extraction to improve reconstruction of human cerebellar connectivity. ACCURAT combines anatomically defined regions, diffusion MRI tractography, and a vertex-level query system that applies fine-grained anatomical constraints at individual points (vertices) along each streamline to extract anatomically consistent pathway segments. These constraints are applied as a post-processing step, enabling ACCURAT to be used with tractography generated by any algorithm. In addition to introducing this framework, we provide a concise, tractography-oriented synthesis of cerebellar pathways grounded in experimental tract-tracing studies. This synthesis organizes pathway-specific origins, trajectories, terminations, decussation patterns, and tractography challenges in a form intended to be accessible and useful to diffusion MRI researchers, thereby informing the design of ACCURAT rules and providing a compact anatomical resource for cerebellar tractography. We then apply ACCURAT using expert-defined anatomical regions and ultra-high-resolution submillimeter diffusion MRI data, enabling detailed reconstruction of multiple cerebellar pathways. We further evaluate the framework across multiple tractography algorithms to assess its generalizability. Together, these contributions establish both an anatomically grounded framework for reconstructing cerebellar connectivity from diffusion MRI and a concise anatomical reference for cerebellar tractography.

## 2. Materials and Methods

### 2.1. Overview of the ACCURAT framework

The ACCURAT framework implements anatomically constrained and curated tractography to reconstruct intrinsic and extrinsic cerebellar pathways from diffusion MRI. An overview of the framework is shown in Figure 1. Diffusion MRI data are used to generate densely seeded whole-brain tractography to ensure adequate coverage of cerebellar pathways for subsequent rule-based reconstruction. Anatomical regions of interest representing nuclei and other structures of the cerebellar connectome are defined from the structural and diffusion MRI data. ACCURAT then performs rule-based post-processing of the tractography using vertex-level queries that evaluate anatomical constraints at individual streamline vertices, extracting anatomically consistent segments.

**Figure 1.**
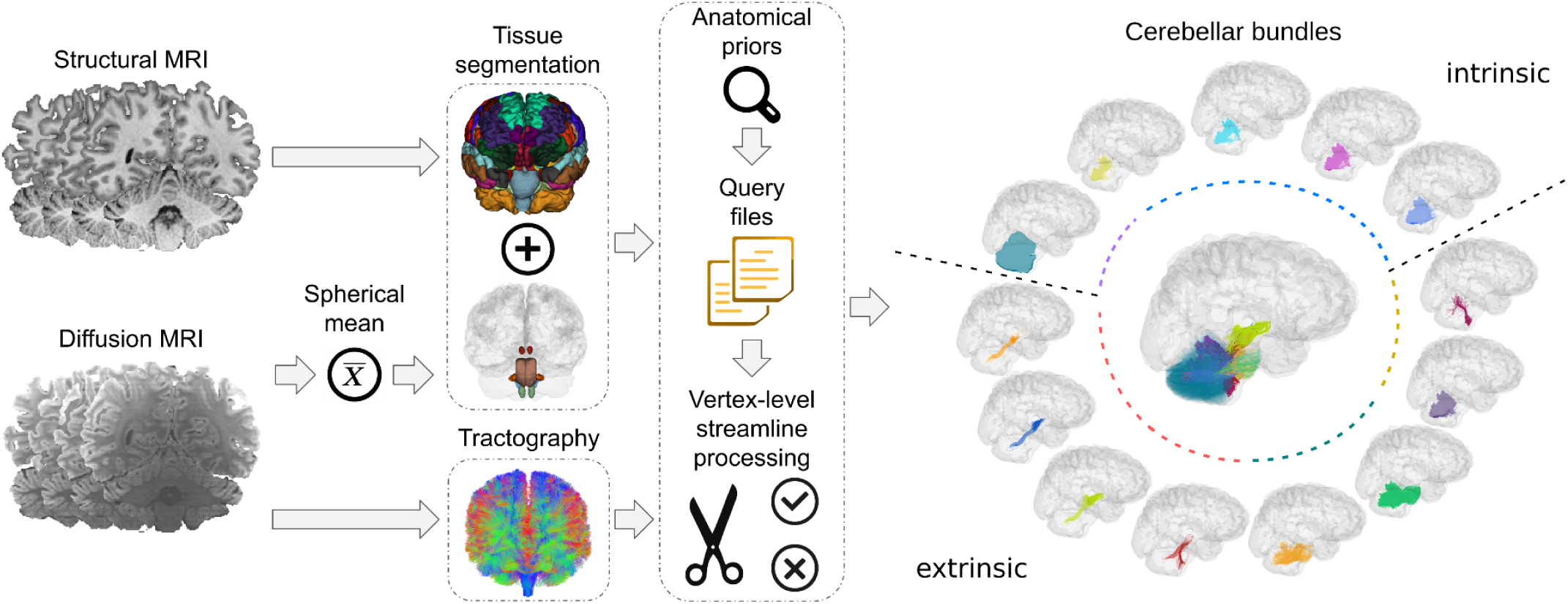
ACCURAT framework. Tissue segmentation from structural MRI (T1w) and diffusion MRI (spherical mean contrast) are used to define anatomical regions of interest. Neuroanatomical priors encoded in query files are then applied to extract bundles of interest from densely seeded diffusion MRI tractography using these ROIs.

The ACCURAT framework is algorithm-agnostic and applicable to results from any tractography method. In this study, we demonstrate its application using both unscented Kalman filter (UKF) tractography (Reddy & Rathi, 2016) and probabilistic parallel transport tractography (PTT) (Aydogan & Shi, 2021). The following sections first describe the ACCURAT pipeline, including its anatomical rules and vertex-level processing, followed by the imaging data, preprocessing, and anatomical segmentation used for pathway reconstruction.

### 2.2. Tract-tracing synthesis and pathway curation

We conducted a structured synthesis of the experimental tract-tracing literature to provide a concise pathway-by-pathway anatomical reference for cerebellar tractography. This synthesis served two purposes: first, to provide an organized summary within the manuscript of cerebellar pathways relevant to tractography; and second, to inform the design of ACCURAT pathway definitions and rules.

Because experimental tract-tracing methods cannot be applied in the human brain, direct evidence of connectional neuroanatomy must be inferred from non-human models (Rushmore, Bouix, et al., 2020). Experimental tract-tracing studies, therefore, served as the foundation for the anatomical pathway definitions used in ACCURAT. The primary source material was tract-tracing research in non-human primates, particularly the macaque monkey. The macaque provides the closest available experimental model in which invasive tract-tracing methods have been applied across multiple components of the pre-cerebellar and cerebellar circuitry. These circuits exhibit a high degree of correspondence across primates, supporting the use of the macaque as a model for human cerebellar connectivity (Jansen & Brodal, 1954; Voogd, 2003). At a gross level, the overall organization of the macaque cerebellum closely parallels that of the human cerebellum and conforms to conserved primate scaling relationships (Herculano-Houzel, 2010). This correspondence, however, is not uniform: the lateral cerebellar hemispheres, particularly crus I and II, show disproportionate expansion in large-brained primates, including humans, consistent with the expansion of prefrontal-cerebellar circuits (Balsters et al., 2010; Luo et al., 2017; MacLeod, 2012; Magielse et al., 2023; Sereno et al., 2020). Where direct tract-tracing evidence in the macaque was incomplete, we drew on information from other animal models, gross anatomical studies, and classic neuroanatomical approaches to complete pathway descriptions.

### 2.3. ACCURAT processing pipeline

ACCURAT reconstructs cerebellar pathways by applying vertex-level streamline processing and neuroanatomically informed rules to tractography data. Unlike previous tractography rule systems that operate on entire streamlines, ACCURAT evaluates anatomical constraints at the level of individual streamline vertices, enabling enforcement of synaptic boundaries within cerebral, brainstem, and cerebellar nuclei.

The ACCURAT framework performs a sequence of anatomically informed operations to isolate cerebellar pathways from densely seeded tractography (Figure 2). The processing pipeline consists of three stages: (1) cortical streamline splitting, which separates segments within the cerebellar cortex from those continuing through cerebellar white matter; (2) vertex-level streamline trimming, which isolates the portion of each trajectory connecting anatomically defined regions while removing segments extending into downstream circuits; and (3) rule-based filtering, which applies inclusion and exclusion constraints derived from cerebellar neuroanatomy. Together, these operations yield candidate streamline reconstructions consistent with known anatomical organization while removing spurious trajectories produced by unconstrained tractography. These steps are described in more detail below.

**Figure 2.**
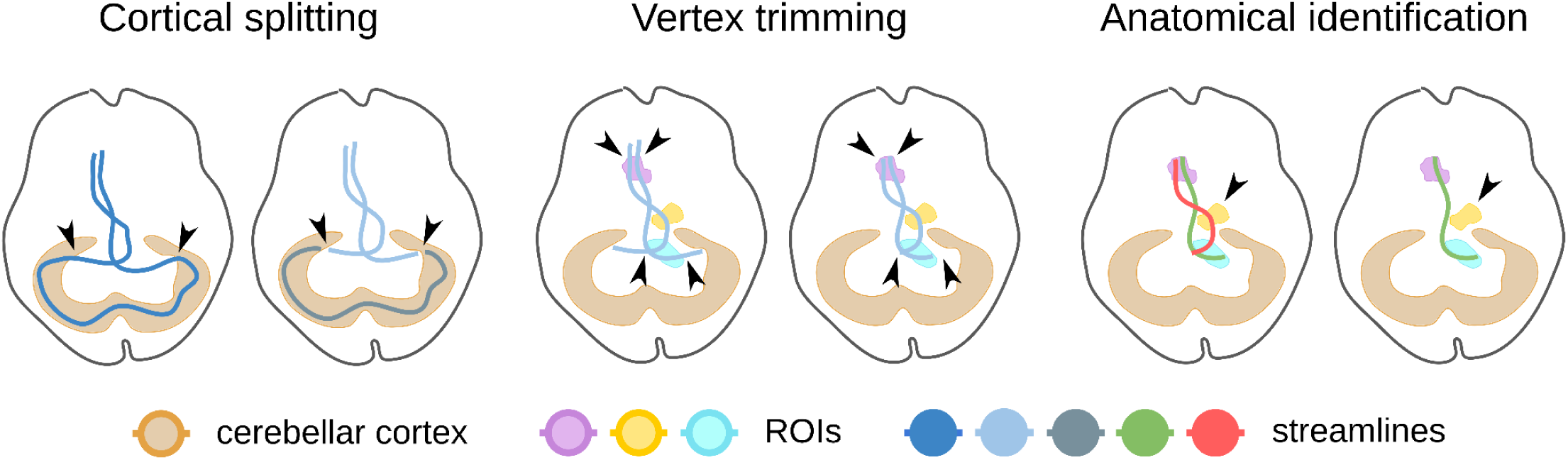
Overview of ACCURAT vertex-level anatomical processing. Cortical streamline splitting, vertex-level trimming at nuclei, and rule-based anatomical identification isolate anatomically consistent pathway segments while removing spurious trajectories. Arrowheads indicate locations where streamlines are cut or removed according to ACCURAT’s processing steps.

#### 2.3.1. Anatomically informed cortical streamline splitting

In ACCURAT, streamline propagation is not restricted to white matter, enabling reconstruction of both intrinsic and extrinsic cerebellar pathways. This design allows tractography to capture trajectories within the cerebellar cortex, including those consistent with the parallel fibers of granule cells (Granziera et al., 2009), which are the most abundant neurons in the mammalian brain (Herculano-Houzel, 2010). Unlike approaches that terminate streamlines at the gray-white matter boundary (e.g., ACT), tractography is allowed to continue through the cerebellar cortex, resulting in streamlines that combine cortical and white matter segments. As an initial processing step, ACCURAT splits these streamlines at the cortical boundary, separating segments confined to the cerebellar cortex from those continuing through cerebellar white matter. Segments within the cerebellar cortex are retained as candidates for the parallel fiber pathway, while remaining segments are used to reconstruct other cerebellar pathways.

#### 2.3.2. Vertex-level streamline trimming at anatomical nuclei

Many cerebellar pathways originate or terminate within specific cerebral, brainstem, or cerebellar nuclei. Because diffusion MRI does not encode synaptic boundaries, tractography streamlines may propagate through these nuclei and continue along downstream pathways, producing trajectories that incorrectly represent multi-synaptic circuits as single anatomical connections.

To address this, ACCURAT performs vertex-level streamline trimming to isolate the portion of each trajectory between anatomically defined regions, thereby restricting endpoints to appropriate nuclei and removing segments that extend into downstream circuits. Streamlines are represented as ordered sequences of vertices, enabling constraints to be evaluated along their trajectory. To enable vertex-level processing, we extend WMQL with a new operator, **`between(region_A, region_B)`**, which trims streamlines to the segment connecting the specified anatomical regions while removing portions of the trajectory that extend beyond either region. The **`between`** operator is symmetric, i.e., **`between(region_A, region_B) = between(region_B, region_A).`**

By operating at the level of individual streamline vertices, this approach enables tractography to respect synaptic endpoints within cerebral, brainstem, and cerebellar nuclei, supporting reconstruction of anatomically defined pathways rather than multi-synaptic circuit trajectories. This also facilitates the investigation of multi-synaptic circuits by enabling their individual anatomical components to be traced separately.

#### 2.3.3. Rule-based identification of cerebellar pathways

The ACCURAT rules define the expected anatomical origins, trajectories, and terminations of cerebellar pathways based on established neuroanatomical knowledge. In particular, they specify the nuclei or cortical regions in which pathways originate or terminate, as well as intermediate anatomical structures that streamlines are required to traverse or avoid.

After vertex-level trimming, candidate segments are evaluated using ACCURAT inclusion and exclusion rules to isolate anatomically plausible trajectories. These rules require streamlines to traverse specific anatomical regions while excluding trajectories entering anatomically implausible structures.

The rules incorporate pathway-specific constraints derived from the neuroanatomical literature. For example, some pathways are required to terminate within deep cerebellar, brainstem, or thalamic nuclei according to established neuroanatomical evidence. Other pathways traverse particular nuclei (e.g., the red nucleus). Similarly, middle cerebellar peduncle pathways, which originate in the pontine nuclei, must pass through the contralateral pontine region before entering the cerebellum.

Applying these rule-based constraints yields candidate cerebellar pathways consistent with known anatomical organization while removing many spurious trajectories produced by unconstrained tractography.

All rules were implemented in the ACCURAT framework as pathway-specific query files to enable reproducible and anatomically consistent reconstruction of cerebellar pathways.

### 2.4. Data acquisition and processing

High-resolution MRI data were acquired from nine healthy adult participants (35 ± 6.5 years old; 5 females / 4 males) on a Siemens Prisma 3T scanner at Brigham and Women’s Hospital (Boston MA, USA). All procedures were approved by the Mass General Brigham Institutional Review Board. Participants received written and verbal information about the purpose and procedures of the imaging experiment. They provided written informed consent prior to imaging.

Structural MRI consisted of a T1-weighted image acquired at 0.7 mm isotropic resolution. Diffusion MRI was acquired using the gSlider sequence (Ramos-Llordén et al., 2020; Setsompop et al., 2018), enabling submillimeter diffusion-weighted imaging as previously described (F. Zhang et al., 2024). The final reconstructed diffusion dataset had 0.76 mm isotropic spatial resolution, providing high spatial detail for mapping cerebellar pathways.

Distortions due to susceptibility, eddy currents, and head motion were corrected using FSL’s *topup* and *eddy* tools (Jenkinson et al., 2012). Because the acquisition included repeated scans, the datasets were first merged to enable motion and eddy-current correction across the full dataset. The repeated acquisitions were then combined using gSlider reconstruction to produce the final diffusion dataset.

The T1-weighted image was processed using FreeSurfer (Fischl, 2012) to generate anatomical segmentations. The T1w image was subsequently co-registered to diffusion space using SynthMorph (Hoffmann et al., 2022, 2024), and the resulting transformation was applied to the FreeSurfer parcellation.

Detailed acquisition parameters and preprocessing procedures are provided in the Supplementary Methods.

Tractography was generated within a brain mask encompassing the cerebellum and extending through associated brainstem and subcortical regions to ensure dense streamline coverage of cerebellar connectivity. As described above, streamline propagation was not restricted to white matter, so that both intrinsic and extrinsic cerebellar pathways could be reconstructed. Two tractography algorithms were used to evaluate the generalizability of the framework: probabilistic parallel transport tractography (PTT) (Aydogan & Shi, 2021) and deterministic multi-tensor unscented Kalman filter tractography (UKF) (Reddy & Rathi, 2016).

For PTT, fiber orientation distributions were estimated using multi-tissue constrained spherical deconvolution with white matter and cerebrospinal fluid response functions obtained using the Dhollander method in MRtrix3 (Tournier et al., 2019). UKF tractography was performed using a two-tensor model. The resulting tractograms were then processed using the same ACCURAT pathway definitions and query rules.

Tractography parameters were selected to provide dense streamline coverage across cerebellar pathways while maintaining anatomically plausible trajectories. Detailed tractography parameters are provided in the Supplementary Methods.

### 2.5. Anatomical regions for cerebellar pathway reconstruction

Anatomical regions of interest were defined to enable rule-based post-processing of tractography streamlines within the ACCURAT framework. These ROIs included cerebellar cortical and white matter structures as well as brainstem and thalamic nuclei involved in cerebellar connectivity. The major anatomical regions used in this study are shown in Figure 3.

**Figure 3.**
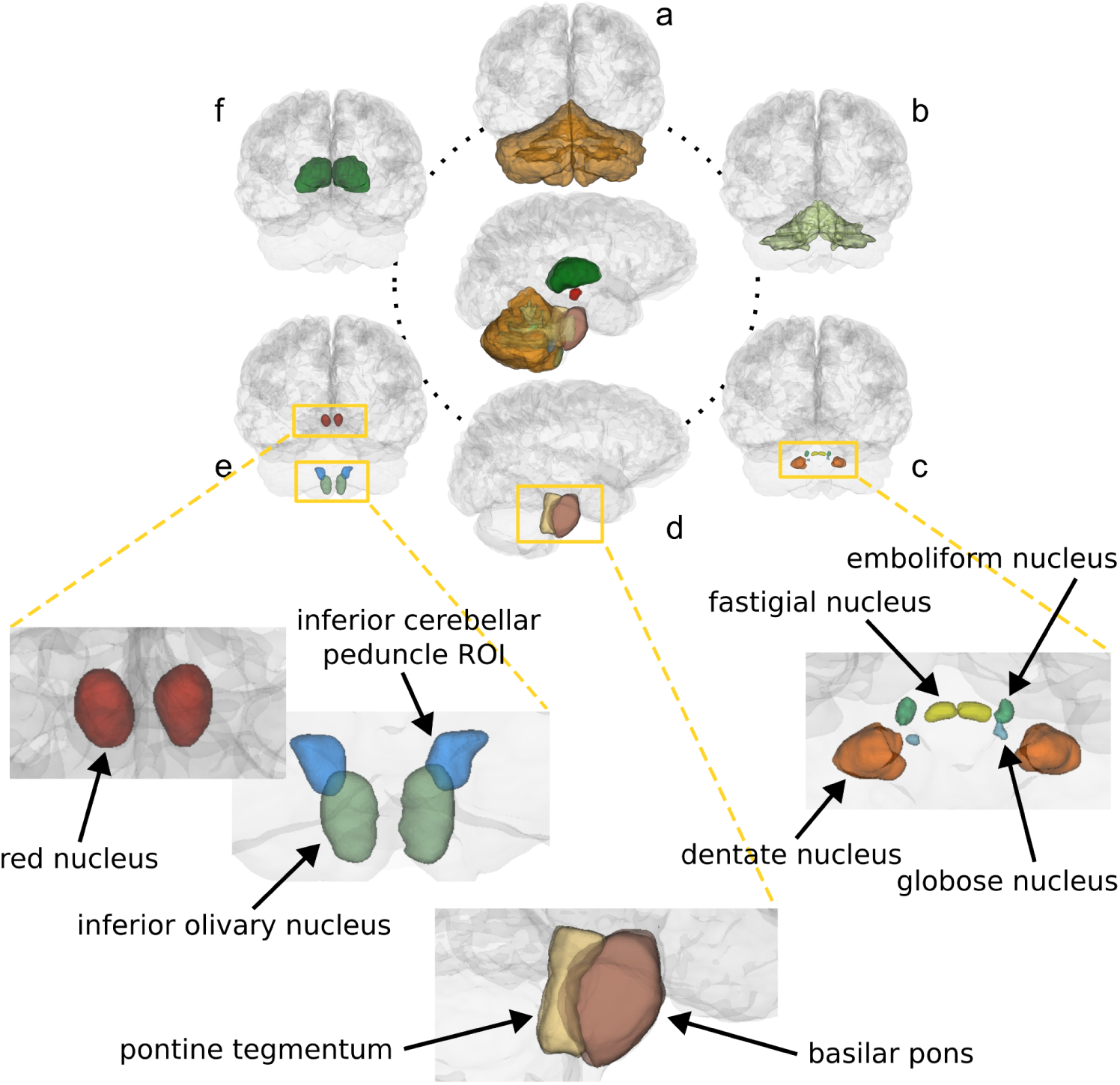
Anatomical regions used for cerebellar tractography. Upper panel: overview of structures included in the reconstruction of cerebellar pathways, including the cerebellar cortex (a), cerebellar white matter (b), deep cerebellar nuclei (c), pontine nuclei (d), inferior olivary nucleus, inferior cerebellar peduncle ROI and red nucleus (e), and thalamus (f). The central circle shows a general anatomical overview. Lower panel: close-up views of selected structures. In this study, the deep cerebellar nuclei (right) include the dentate, emboliform, globose, and fastigial nuclei. The pontine nuclei (middle) are subdivided into two adjacent regions: the basilar pons and the pontine tegmentum. Additional close-up views (left) show the inferior olivary nucleus, inferior cerebellar peduncle ROI, and red nucleus. All structures are segmented bilaterally.

#### 2.5.1. Automated segmentation

Automated anatomical segmentation was performed using FreeSurfer (Fischl, 2012). These segmentations provided masks of the cerebellar cortex, cerebellar white matter, and thalamus, together with additional anatomical regions used as exclusion ROIs to remove anatomically implausible streamlines.

#### 2.5.2. Anatomical curation of tractography constraints

ACCURAT reconstructs pathways using anatomically defined constraints that specify the expected origins, trajectories, and terminations of axons. Implementing these rules requires anatomical regions that can be reliably delineated in MRI.

While automated methods have been proposed for dMRI-based segmentation of larger deep cerebellar nuclei (primarily dentate and interposed, which combines globose and emboliform (Gaviraghi et al., 2021; Kim et al., 2020; Legarreta et al., 2025; Noguera et al., 2019)), these approaches are not designed for submillimeter dMRI acquisitions and do not provide all regions necessary for ACCURAT. Therefore, we performed a neuroanatomical evaluation to determine which cerebellar and brainstem structures could be reliably identified using spherical mean diffusion contrast.

Using a diffusion-derived contrast enables anatomical delineation directly in diffusion MRI space, avoiding geometric distortions and inter-modality registration errors that can occur when T1-based segmentations are mapped into diffusion space (X. Wang et al., 2026; F. Zhang et al., 2021). These errors disproportionately affect small structures such as cerebellar and brainstem nuclei. The spherical mean provides a stable diffusion-derived contrast that enhances visualization of these structures and has been shown to improve cerebellar segmentation performance relative to other diffusion MRI contrasts (Legarreta et al., 2025).

Based on this evaluation, segmentation feasibility was determined by evaluating whether individual nuclei could be consistently identified based on contrast and anatomical boundaries. Several nuclei involved in cerebellar connectivity could be reliably segmented, including the inferior olivary nucleus, red nucleus, and deep cerebellar nuclei. Other nuclei were either too small, lacked sufficient contrast, or were not anatomically distinguishable as discrete structures on MRI. In some cases, ACCURAT rules could be defined using anatomically meaningful bounding regions that encompassed the underlying nuclei. For example, pontocerebellar projections arise from multiple pontine nuclei that are not individually visible in MRI; therefore, the pons (including pontine tegmentum and basilar pons) was segmented as a bounding region representing their origin.

This anatomically informed strategy enables systematic definition of tractography constraints grounded in structures that are reliably identifiable in in vivo human data, while maintaining consistency with known neuroanatomy. Accordingly, only pathways whose source and target regions (or bounding regions) could be reliably delineated were included in the ACCURAT framework. Here, “source” and “target” refer to known neuroanatomical origins and terminations; however, because diffusion MRI tractography does not encode axonal directionality, reconstructed streamlines should be interpreted as undirected trajectories constrained by these regions.

#### 2.5.3. Expert neuroanatomical segmentation

Expert neuroanatomical segmentation was performed on the spherical mean diffusion image by a team with extensive expertise in brainstem and cerebellar anatomy (RJR, NM, and EY) (Makris et al., 2003, 2005; Rushmore, Wilson-Braun, et al., 2020; Zheng et al., 2023). Using this contrast, key nuclei and brainstem regions relevant for cerebellar pathway reconstruction were manually delineated, including the deep cerebellar nuclei (dentate, emboliform, globose, and fastigial), the inferior olivary nucleus, the red nucleus, subdivisions of the pontine region (basilar pons and pontine tegmentum), and a region through which the inferior cerebellar peduncle passes. To perform segmentation of these structures, an intensity-based thresholding tool (3D Slicer; (Fedorov et al., 2012)) was first used to delineate borders, followed by expert manual refinement and editing based on known anatomical boundaries.

All FreeSurfer-derived and expert-defined ROIs were merged into a single labelmap for each participant. These regions were then used to define anatomical constraints for tractography within the ACCURAT framework.

## 3. Results

Expert neuroanatomical segmentation was successfully performed (Table S1 provides volumes of the segmented structures), enabling the derivation of ACCURAT rules for multiple extrinsic and intrinsic cerebellar pathways.

### 3.1. Reconstruction of cerebellar pathways with ACCURAT

#### 3.1.1. Extrinsic cerebellar pathways

The extrinsic cerebellar connections are conveyed through the three cerebellar peduncles—the inferior, middle, and superior peduncles—each of which contains multiple distinct anatomical pathways. These include both afferent (input) pathways that convey signals to the cerebellum and efferent (output) pathways that carry cerebellar signals to other regions of the nervous system. In addition, we describe a smaller extrinsic pathway, the uncinate fasciculus of the cerebellum, which hooks over the superior cerebellar peduncle.

##### Inferior cerebellar peduncles (ICP)

###### Neuroanatomical organization

The inferior cerebellar peduncle (ICP) is composed of two divisions: the lateral division, the restiform body, which carries the major afferent fiber systems, and the medial division, the juxtarestiform body, which carries fibers interconnecting the cerebellum with the vestibular nuclei and other brainstem targets. The ICP contains multiple afferent pathways conveying proprioceptive input from ipsilateral musculature and error-signaling input from the contralateral inferior olivary nucleus, as well as axons that interconnect the cerebellum with the brainstem vestibular system and other brainstem nuclei. The ICP also carries efferent fastigial projections to brainstem nuclei via the juxtarestiform body, including fastigiovestibular (Asanuma et al., 1983b; Batton et al., 1977; Kalil, 1979) and fastigiotectal (Batton et al., 1977; May et al., 1990; Yamada & Noda, 1987) tract axons. In addition, the ICP includes Purkinje cell axons that project directly to the vestibular nuclei, as well as vestibulo-cerebellar axons (A. Brodal & Brodal, 1985; Nagao et al., 1997a, 1997b; Noda et al., 1990).

Proprioceptive signals are conveyed via the dorsal (posterior) spinocerebellar tract, which carries input from the ipsilateral lower body; the cuneocerebellar tract, which carries input from the ipsilateral upper body; and the trigeminocerebellar tract, which carries input from the ipsilateral muscles of the head (Jansen & Brodal, 1954; Larsell & Jansen, 1972; M. C. Smith, 1957). These pathways are composed of mossy fibers, which provide indirect input to Purkinje cells via granule cells in the cerebellar cortex. The mossy fiber axons also send branches, termed collaterals, to the deep cerebellar nuclei. Some fibers from the reticular formation, primarily from the paramedian reticular nucleus, also travel in the ICP to the cerebellum (Somana & Walberg, 1978).

A major contributor to the ICP is the olivocerebellar projection, involved in error signaling, which arises from the contralateral inferior olivary nucleus and provides climbing fiber input that synapses directly on Purkinje cells. These axons also give off collateral branches to the deep cerebellar nuclei. The inferior olivary nucleus, which can be segmented in our data, is therefore an especially important anatomical landmark for reconstructing this component of the ICP. The course of olivocerebellar fibers is well documented: these axons cross the midline and pass superior to, and in some cases through, the contralateral inferior olivary nucleus before entering the ICP (P. Brodal & Brodal, 1981; Ikeda et al., 1989) Their collateral branches to the deep cerebellar nuclei are smaller-caliber and potentially unmyelinated branches of the parent axons (Groenewegen et al., 1979; Matsushita & Ikeda, 1970), which then ramify to form synaptic terminals.

The major pathways contributing to the ICP and their anatomical characteristics are summarized in Table 1.

**Table 1.** Major pathways contributing to the inferior cerebellar peduncle (ICP). Though quantitative data on their exact axon counts are lacking in human and non-human primates, an asterisk (*) denotes bundles generally believed to contribute the most axons (olivocerebellar, spinocerebellar, and cuneocerebellar pathways). Feasibility of expert neuroanatomical segmentation of nuclei is indicated in italics within parentheses.

| Pathway | Fiber type | Input / Output | Source | Target | Decussation | ACCURAT implementation |
| --- | --- | --- | --- | --- | --- | --- |
| **Inferior cerebellar peduncle (ICP)** | Mixed | Mixed | ICP white matter region (*segmentable*) | Cerebellar cortex | Yes/No | Peduncle reconstruction |
| *Restiform body* |  |  |  |  |  |  |
| **Olivocerebellar \*** | Climbing | Input | Inferior olivary nucleus (*segmentable*) | Cerebellar cortex | Yes | Pathway reconstruction |
| Dorsal (posterior) spinocerebellar * | Mossy | Input | Nucleus dorsalis of Clarke of spinal cord (*outside acquired data*) | Cerebellar cortex | No | — |
| Cuneocerebellar* | Mossy | Input | External cuneate nucleus (*difficult*) | Cerebellar cortex | No | — |
| Medullary reticulocerebellar | Mossy | Input | Medullary reticular formation nuclei (*difficult*) | Cerebellar cortex | No | — |
| Trigemino-cerebellar | Mossy | Input | Spinal nucleus of trigeminal, primary sensory nucleus of trigeminal (*difficult*) | Cerebellar cortex | No | — |
| *Juxtarestiform body* |  |  |  |  |  |  |
| Vestibulocerebellar | Mossy | Input | Vestibular nuclei, Vestibular (Scarpa's) ganglion (*difficult*) | Cerebellar cortex | No | — |
| Fastigiovestibular | N/A | Output | Fastigial nucleus (*segmentable*) | Vestibular nuclei (*difficult*) | No | — |
| Fastigiotectal | N/A | Output | Fastigial nucleus (*segmentable*) | Superior Colliculus (*segmentable*) | Yes (cross in cerebellum) | — |
| Purkinje cortico-vestibular | Purkinje | Output | Cerebellar cortex | Vestibular nuclei (*difficult*) | No | — |

###### Tractography challenges

Reconstruction of ICP pathways is complicated by the presence of multiple small nuclei that cannot be reliably segmented in MRI (Table 1), limiting the ability to delineate many ICP subcomponents individually using diffusion MRI. Given this challenge, ACCURAT focuses on pathways whose key nuclei can be reliably identified, such as the olivocerebellar pathway. The olivocerebellar pathway is particularly difficult to trace (Granziera et al., 2009; Yin et al., 2023) due to its challenging trajectory, including a decussation and traversal of complex brainstem regions with crossing fibers and multiple nuclei. Although the fastigial nucleus and superior colliculus were also visible in our data, we did not attempt to trace the small fastigiotectal bundle as its trajectory is complex: it originates in the fastigial nucleus, crosses the cerebellar midline, traverses the contralateral fastigial nucleus, and then travels via the juxtarestiform portion of the ICP to the superior colliculus (Batton et al., 1977; May et al., 1990).

These anatomical considerations inform the design of ACCURAT rules for ICP reconstruction.

###### ACCURAT rule — olivocerebellar pathway

The inferior olivary nucleus could be reliably segmented in the spherical mean data. Thus, ACCURAT identifies candidate olivocerebellar trajectories between the inferior olivary nucleus and the contralateral cerebellar cortex. Additional exclusion criteria remove spurious streamlines entering the pons, ipsilateral cerebellar white matter, or other regions such as cerebral structures. An example ACCURAT query implementing these constraints is shown in Box 1; the complete set of pathway queries is available in the ACCURAT repository (see *Data and Code Availability*).

###### Box 1. Example illustrative ACCURAT natural-language query.

~~~
olivocerebellar.side = (
   between(inferior_olive.side, cerebellum_cortex.opposite)
   not in pons
   not in cerebellum_cortex.side
   not in cerebellum_white_matter.side
   *<ADDITIONAL exclusion ROIs for spurious streamline control>*
)
~~~

###### ACCURAT rule — ICP

Because many contributing nuclei could not be reliably segmented or were outside the imaging field of view (Table 1), an additional ROI positioned at the dorsolateral aspect of the medulla was introduced to reconstruct the ICP as a whole. This ROI captures the major fiber bundle forming the peduncle, even when individual constituent pathways cannot be reliably separated. The ICP ACCURAT rule thus identifies candidate trajectories between the ICP ROI and the ipsilateral cerebellar cortex.

##### Resulting reconstructions

Using these constraints, ACCURAT yields trajectories consistent with known olivocerebellar pathways and inferior cerebellar peduncles (Figure 4). Although reconstruction of the olivocerebellar pathway was feasible (Figure 4, top row), it remained challenging and was not identified in all subjects (Table 4).

**Figure 4.**
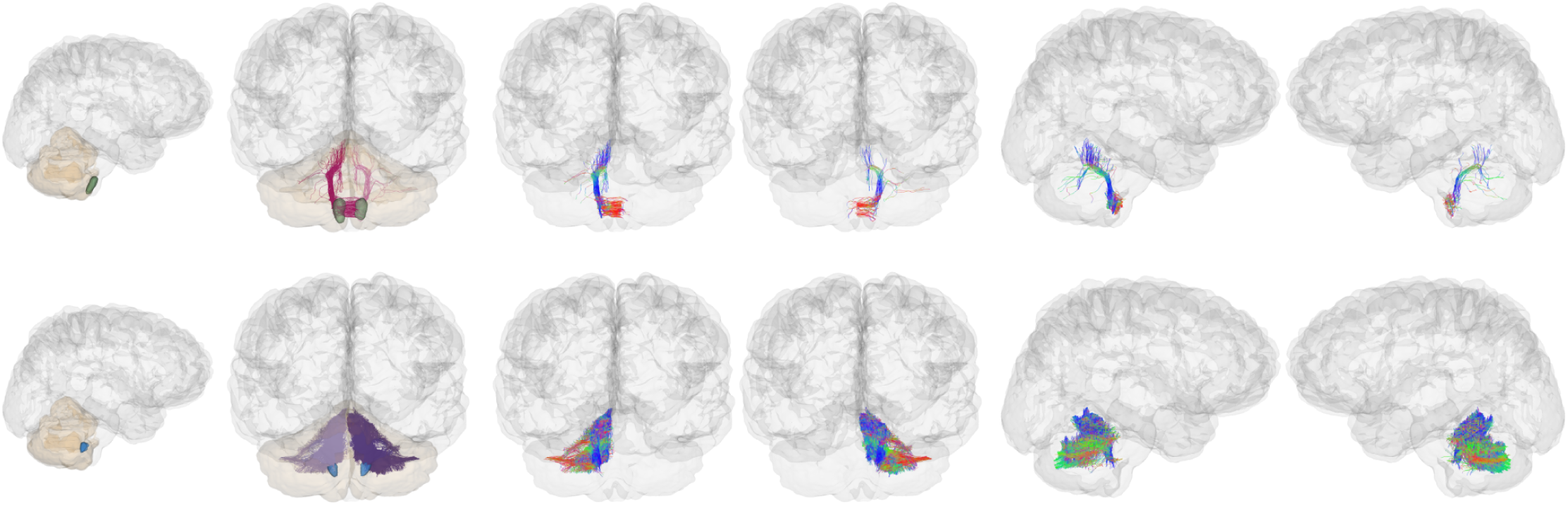
Inferior cerebellar peduncles as extracted by ACCURAT in an exemplar participant, using PTT data. Top row: olivocerebellar pathway; bottom row: inferior cerebellar peduncle. The source and target regions of interest are shown in the leftmost column.

##### Middle cerebellar peduncles (MCP)

###### Neuroanatomical organization

The middle cerebellar peduncles, also known as the brachia pontis, are two large fiber bundles located on the lateral aspects of the brainstem. The axons within each MCP project to the cerebellum as mossy fibers, which synapse with neurons of the granule cell layer of the cerebellar cortex. Many of these mossy fiber axons also give rise to collateral branches to the deep cerebellar nuclei.

Pontocerebellar projections are the dominant component of the MCP and arise from neurons in the pontine nuclei, within the basilar (ventral) pons. The vast majority of pontocerebellar fibers cross the midline before entering the contralateral cerebellum: pathway tracing studies from the macaque indicate that after unilateral injection of retrograde tracer in the lateral cerebellar hemispheres, 90% of neurons in the pontine nuclei are labeled on the contralateral side, whereas this number decreases to 70% after vermal injection (P. Brodal, 1979, 1982).

The MCP also contains projections from other brainstem inputs to the cerebellum, including axons from the reticular formation, particularly the nucleus reticularis tegmenti pontis (NRTP). The majority of these reticulocerebellar projections (approximately 64%) are to the contralateral cerebellum and specifically to the vermal and flocculonodular regions (P. Brodal, 1980, 1982; Langer et al., 1985).

A topographic organization of the MCP has been described, in which axons from the lateral, ventral, and rostral pontine nuclei occupy more ventral (anterior) portions of the peduncle, whereas axons from the NRTP and more caudal pontine nuclei occupy more dorsal (posterior) regions. The principal anatomical pathways contributing to the middle cerebellar peduncles are summarized in Table 2.

**Table 2.** Major pathways contributing to the middle cerebellar peduncles (MCP). An asterisk (*) denotes the bundle(s) containing the largest number of axons.

| Pathway | Fiber type | Input / Output | Source | Target | Decussation | ACCURAT implementation |
| --- | --- | --- | --- | --- | --- | --- |
| **Pontocerebellar \*** | Mossy | Input | Pontine nuclei region (*forms the majority of gray matter in the basilar pons*) | Cerebellar cortex (widespread; ~3 times denser projections to lateral cerebellar hemispheres) | Yes | Pathway reconstruction |
| **Pontine reticulocerebellar** | Mossy | Input | Nucleus reticularis tegmenti pontis (*not visible; included within pontine tegmentum ROI*) | Cerebellar cortex (primarily vermal and flocculonodular) | Yes | Pathway reconstruction |

###### Tractography challenges

Without explicit anatomical constraints in the pontine region, tractography frequently reconstructs spurious MCP trajectories connecting the left and right cerebellar cortices (Warrington et al., 2022; F.-C. Yeh et al., 2018; F. Zhang et al., 2018), despite the absence of known direct anatomical connections between these regions. Other approaches address this by selecting streamlines that connect the pons to the contralateral cerebellum while excluding trajectories entering the ipsilateral cerebellum, thereby capturing the expected decussation of the MCP (Martinez et al., 2026). However, this whole-streamline selection strategy relies on streamlines that happen to terminate in the pons—an outcome more common in DTI tractography (Martinez et al., 2026) using, for example, strict FA stopping thresholds, but much less likely in probabilistic or higher-order models, where anatomical rules are often needed for streamline termination. Furthermore, we do not attempt to trace minor uncrossed MCP projections or collateral branches to deep cerebellar nuclei.

###### ACCURAT rules — MCP

ACCURAT identifies candidate pontocerebellar and reticulocerebellar trajectories between the pons (using the basilar pons and pontine tegmentum ROIs) and the contralateral cerebellar cortex. Reconstructed streamlines are required to pass through the contralateral pons, consistent with the primarily decussating or crossed neuroanatomy of these MCP components. Additional exclusion criteria remove streamlines entering spurious brainstem, cerebrum, or ventricular areas.

##### Resulting reconstructions

These constraints, integrated within the ACCURAT framework, yield trajectories consistent with the known anatomy of pontocerebellar and pontine reticulocerebellar projections within the middle cerebellar peduncle (Figure 5).

**Figure 5.**
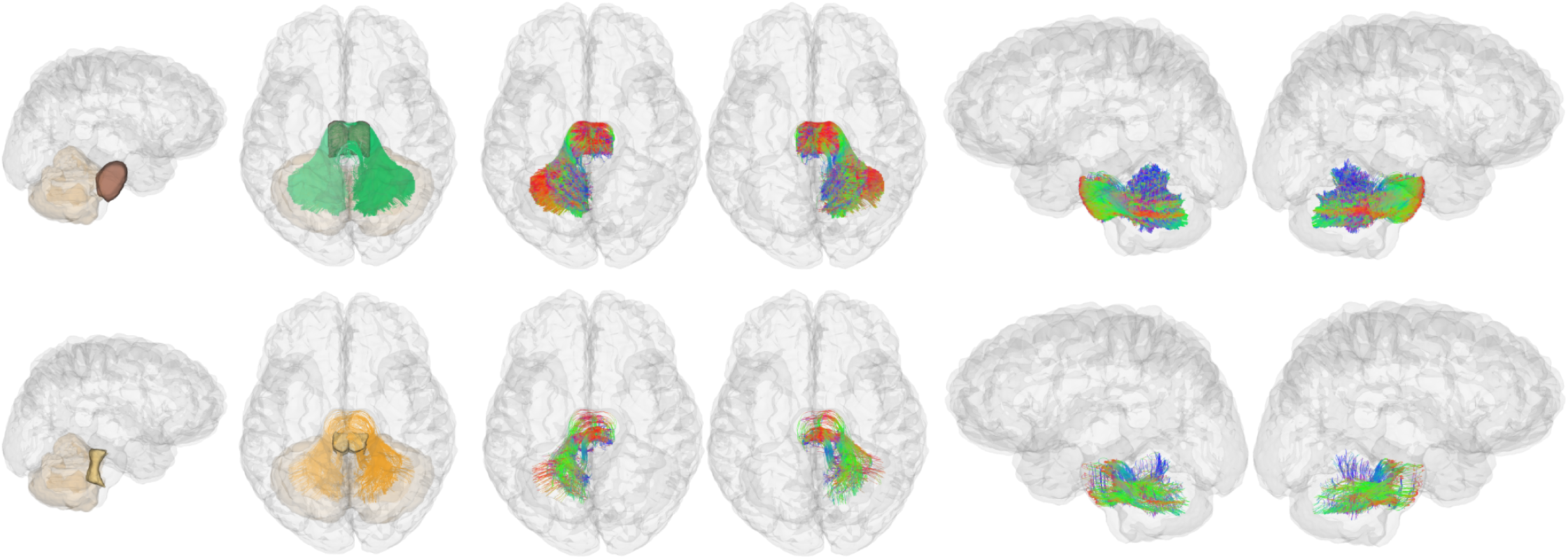
Middle cerebellar peduncles as extracted by ACCURAT in an exemplar participant, using PTT data. Top row: pontocerebellar (from basilar pons); bottom row: pontine reticulocerebellar (from pontine tegmentum).

##### Superior cerebellar peduncles (SCP)

###### Neuroanatomical organization

The superior cerebellar peduncle is the major output pathway of the cerebellum and primarily carries axons originating from the dentate, globose, and emboliform nuclei. Most SCP fibers travel anteriorly into the brainstem tegmentum, decussate, and then travel superiorly to terminate in the contralateral red nucleus and thalamus.

Dentate nucleus outputs dominate SCP projections. The axons of the dentate nucleus project predominantly to the parvocellular (rostral) portion of the red nucleus and to the ventral lateral nucleus of the thalamus, with additional projections to the ventral anterior, central lateral, and ventral posterior lateral thalamic nuclei (Amino et al., 2001; Asanuma et al., 1983a, 1983b, 1983c; Bostan & Strick, 2013; Chan-Palay, 1977; Gonzalo-Ruiz et al., 1988; Hoshi et al., 2005; Kalil, 1979; Kyuhou et al., 1997; Mason et al., 2000; Sakai et al., 1996). Projections from the globose and emboliform nuclei have more limited thalamic projections and terminate primarily in the contralateral red nucleus.

In addition to the projections to the red nucleus and the thalamus, other efferent cerebellar projections cross in the decussation of the SCP and project to the contralateral inferior olive (including dentato-olivary and interposed-olivary pathways (Asanuma et al., 1983b; Ilinsky & Kultas-Ilinsky, 1984; Kalil, 1979) and nuclei of the reticular formation (Asanuma et al., 1983b; Kalil, 1979). The SCP also contains projections that project to the superior colliculus and smaller nuclei related to eye movements, some of which stay ipsilaterally, and others that cross in the brainstem (Batton et al., 1977; May et al., 1990, 1992). The SCP also carries reciprocal connections between the cerebellum and hypothalamus (Haines et al., 1990). The SCP also carries the ventral (anterior) spinocerebellar tract, a small afferent pathway from spinal cord border neurons that crosses at the spinal cord level and recrosses within the cerebellar white matter (Jansen & Brodal, 1954).

The principal anatomical pathways contributing to the superior cerebellar peduncle are summarized in Table 3.

**Table 3.** Major pathways contributing to the superior cerebellar peduncles (SCP). An asterisk (*) denotes the bundle(s) containing the largest number of axons (dentate nucleus projections).

| Pathway | Input / Output | Ascending / Descending | Source nucleus | Target | Decussation | ACCURAT implementation |
| --- | --- | --- | --- | --- | --- | --- |
| **Dentato-rubral fibers \*** | Output | Ascend | Dentate nucleus<br>(*segmentable*) | Red nucleus<br>(*segmentable*) | Yes | Pathway reconstruction |
| **Dentato-thalamic \*** | Output | Ascend | Dentate nucleus<br>(*segmentable*) | Ventral lateral thalamus<br>(*segmentable*) | Yes | Pathway reconstruction |
| **Dentato-rubro-thalamic \*** | Output | Ascend | Dentate nucleus<br>(*segmentable*) | Red nucleus and thalamus<br>(*segmentable*) | Yes | Pathway reconstruction |
| **Dentato-olivary** | Output | Descend | Dentate nucleus<br>(*segmentable*) | Inferior olive<br>(*segmentable*) | Yes | Pathway reconstruction |
| Interposed-olivary | Output | Descend | Globose + emboliform nuclei<br>(*segmentable*) | Inferior olive<br>(*segmentable*) | Yes | *Candidate pathway* (definition provided) |
| Interposed-rubral | Output | Ascend | Globose + emboliform nuclei<br>(*segmentable*) | Red nucleus<br>(*segmentable*) | Yes | *Candidate pathway* (definition provided) |
| Interposed-thalamic | Output | Ascend | Globose + emboliform nuclei<br>(*segmentable*) | Thalamus<br>(*segmentable*) | Yes | *Candidate pathway* (definition provided) |
| Ventral (anterior) spinocerebellar tract | Input |  | Spinal cord border neurons<br>(*not segmentable*) | Cerebellar Cortex<br>(*segmentable*) | Crosses at spinal cord, then again in cerebellar white matter | — |
| Cerebello-hypothalamic | Output | Ascend | Globose, Emboliform, Dentate (*segmentable*) | Lateral and Posterior hypothalamic nuclei (*segmentable*) | Yes; some axons cross for a second time in diencephalon | — |
| Hypothalamo-cerebellar | Input | Descend | Hypothalamic nuclei (lateral and posterior hypothalamic nuclei, lateral mammillary and supramammillary nuclei) (*segmentable but difficult*) | Cerebellar Cortex (*segmentable*) | Yes (crossing location unknown) | — |
| Cerebello-tectal | Output | Ascend | Globose, Dentate (*segmentable*) | Superior Colliculus(*segmentable*) | Yes, some axons cross for a second time in brainstem | — |
| Cerebello-reticular | Output | Descend | Globose, Emboliform, Dentate (*segmentable*) | Reticular Formation (*segmentable but difficult*) | Yes | — |

**Table 4.** Minor efferent pathway through the uncinate fasciculus of the cerebellum.

| Pathway | Input / Output | Source nucleus | Target | Decussation | ACCURAT implementation |
| --- | --- | --- | --- | --- | --- |
| Various fastigial brainstem projections | Output | Fastigial nucleus<br>(*segmentable*) | Various brainstem nuclei<br>(*difficult*) | Yes (within cerebellum) | — |
| Fastigio-thalamic | Output | Fastigial nucleus<br>(*segmentable*) | Thalamus | Yes (within cerebellum) | — |

###### Neuroanatomical note on SCP decussation

In the classic neuroanatomical literature, the SCP axons are described as fully decussating at the level of the caudal midbrain (Asanuma et al., 1983b; Carpenter & Stevens, 1957; Carpenter & Sutin, 1983; Jansen & Brodal, 1954; Kalil, 1979; Larsell & Jansen, 1972; Nieuwenhuys et al., 2008; ten Donkelaar et al., 2020). The completeness of the SCP decussation is also observed in most macaque experimental neuroanatomical reports using tract tracers (Asanuma et al., 1983b; Kalil, 1979). However, recent data from diffusion MRI, microdissection, and physiological recordings in the human brain suggest that a substantial number of cerebellothalamic fibers remain uncrossed (Brogna et al., 2022; Meola, Comert, et al., 2016; Petersen et al., 2018; Sajonz et al., 2024). Non-decussating fibers have been sparsely labeled in feline tracer studies (Ilinsky et al., 1987; Nakano et al., 1980; Sugimoto et al., 1981), and in non-human primates the evidence is restricted to a single report of anterograde tracing in one macaque (Chan-Palay, 1977). In that study, a small number of labeled fibers in the SCP from the posterior dentate nucleus did not decussate at the midbrain but instead ascended ipsilaterally to the thalamus, where they crossed in the interthalamic adhesion. Taken together, experimental tracing studies in animal models, including the macaque, provide limited support for a substantial population of non-decussating SCP fibers. However, findings from the human brain raise the possibility that this uncrossed pathway may be more prominent or potentially expanded in the human brain. This interpretation should be tempered by the known limitations of dMRI tractography, which performs poorly in regions of complex fiber architecture and crossings, such as decussations.

###### Tractography challenges

Reconstruction of the SCP decussation is particularly challenging because of the difficulty of tracing crossing fibers, which still affects modern multi-fiber tractography approaches (Coenen et al., 2021). This limitation biases tractography towards reconstruction of ipsilateral streamlines and likely contributes to the prevalence of non-decussating SCP streamlines reported in previous studies. Isolating SCP projections is also difficult because streamlines may continue propagation through cerebellar nuclei (Tchetchenian et al., 2024) or continue through the thalamus into cerebral white matter in the absence of anatomical constraints (Habas & Manto, 2018). Most tractography studies, therefore, focus on the dominant dentate projections, commonly referred to as dentato-rubro-thalamic, dentatothalamic, dentato-rubro-thalamo-cortical, DRT, or DRTT projections, depending on the trajectory traced. The descending fibers of the SCP have received much less attention in the tractography literature. For example, the dentato-rubro-olivary pathway (actually comprising two pathways, dentato-rubral and rubro-olivary, with a synapse in the red nucleus) has been reconstructed at 7T (Calzoni et al., 2025), whereas the direct dentato-olivary pathway, which turns below the red nucleus to descend to the contralateral inferior olive, has received little prior attention to our knowledge. While we included the dentato-rubral pathway here, we did not include the rubro-olivary pathway, as it is located in the brainstem, and this work focuses on cerebellar pathways specifically. Tracing the cerebello-hypothalamic pathways is challenging, in part due to their small size, and has been attempted with limited success (Çavdar et al., 2020). We did not attempt to trace doubly decussating pathways, such as parts of the cerebello-hypothalamic and cerebello-tectal pathways, or pathways that cross between cerebellar hemispheres in the white matter, such as the ventral spinocerebellar pathway. Finally, precise determination of streamline endpoints within small source nuclei is challenging due to their complex morphology; for example, the dentate nucleus consists of a highly folded, pleated sheet that is only coarsely represented by the convex structure visible in most MRI acquisitions.

###### ACCURAT rules — Ascending SCP

ACCURAT identifies candidate ascending trajectories between the deep cerebellar nuclei and the contralateral red nucleus and thalamus as relevant for each pathway. Further exclusion criteria remove spurious streamlines entering anatomically inconsistent regions such as ventricles, contralateral pontine regions, or cerebral regions.

###### ACCURAT rules — Descending SCP

The dentato-olivary pathway is extracted by ACCURAT between the dentate nucleus and the contralateral olivary nucleus, excluding ascending trajectories (e.g., toward the red nucleus or thalamus), or artifactual connections traversing the cerebellar midline. To exclude olivocerebellar pathways, which travel through the ICP and send collaterals to the dentate, the ipsilateral ICP ROI is employed as an exclusion ROI.

##### Resulting reconstructions

Within ACCURAT, these constraints enable the extraction of the decussating trajectories of the superior cerebellar peduncle bundles (Figure 6).

**Figure 6.**
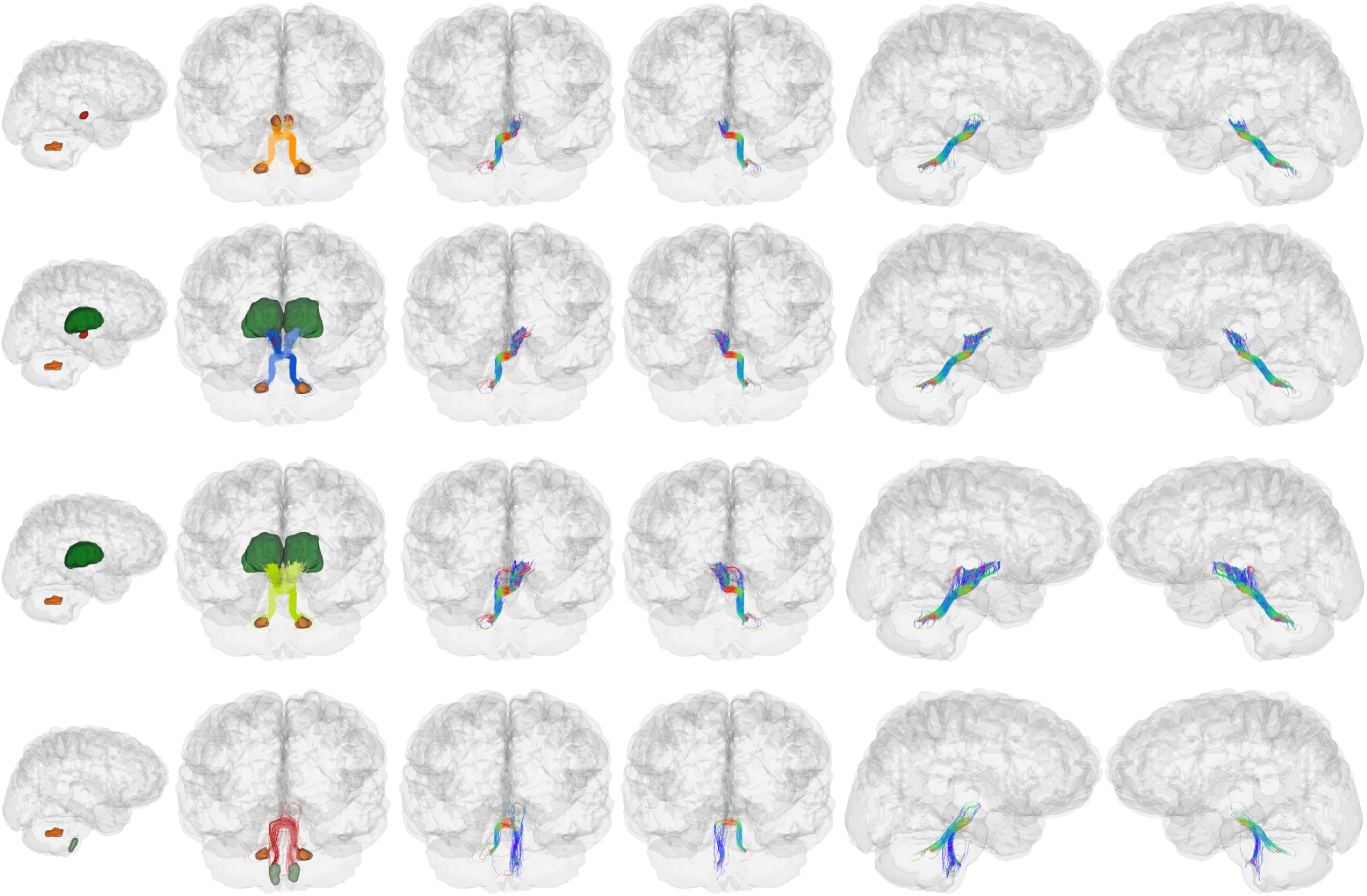
Superior cerebellar peduncles as extracted by ACCURAT in an exemplar participant, using PTT data. Top row: dentato-rubral; second row: dentato-rubro-thalamic; third row: dentato-thalamic; bottom row: dentato-olivary.

##### Fastigial projections: the uncinate fasciculus of the cerebellum

In addition to the peduncles, we describe here for completeness the extrinsic projections of the fastigial nucleus, an additional efferent (cerebellar output) pathway that is less well known.

###### Neuroanatomical organization

In addition to the extrinsic cerebellar connections conveyed through the cerebellar peduncles, efferent projections from the fastigial nucleus reach contralateral brainstem nuclei (and diencephalic nuclei) through the uncinate fasciculus of the cerebellum (Asanuma et al., 1983b; Batton et al., 1977; Ikeda et al., 1989; Kalil, 1979; Noda et al., 1990; Sato & Noda, 1991), also known as the hook bundle of Russell. This bundle constitutes the primary efferent route for axons originating in the fastigial nucleus. Axons arising from the fastigial nucleus cross the midline within cerebellar white matter, may pass over or through the contralateral fastigial nucleus, wrap around the superior and lateral borders of the superior cerebellar peduncle within the cerebellum, and then turn to reach brainstem targets (Table 4). This fasciculus should not be confused with the uncinate fasciculus of the cerebrum, a major association pathway connecting the frontal and temporal lobes. The cerebellar uncinate fasciculus described here forms part of the extrinsic cerebellar connectome.

###### Tractography challenges

Although we note this pathway here for completeness, its small size, its course through and around the contralateral fastigial nucleus, its proximity to the superior cerebellar peduncle, and the difficulty of segmenting many of the relevant brainstem nuclei have thus far precluded its investigation with diffusion MRI, to our knowledge, including in the present study. In addition to its small size, this pathway involves multiple tractography reconstruction challenges, including cerebellar midline crossing, potential traversal of a deep cerebellar nucleus, and a sharp turn in trajectory.

##### A note on collaterals in the cerebellum

Collaterals are a ubiquitous feature of both cerebral and cerebellar circuits. In general, collaterals are branches of an axon. In the cerebral cortex, they can be divided into intrinsic collaterals, which arise near the cell body and terminate on nearby neurons, and extrinsic collaterals, which branch distally along the axon and contact targets en route to their main target (Rockland, 2013, 2018; Rockland & Rushmore, 2025). Collateral branches are typically of smaller caliber and may be unmyelinated. Most recent research on collaterals has been done in the cerebrum, where collaterals are common in long-range cortical projection systems (Coudé et al., 2018; Kita & Kita, 2012; Rockland & Rushmore, 2025).

In the cerebellum, collaterals are an important part of signal flow and occur in most pathways. Both mossy fibers and climbing fibers (see ICP and MCP sections, above) end in the cerebellar cortex, but along the way they send collateral branches to the deep cerebellar nuclei. In this way, the deep cerebellar nuclei receive inputs prior to processing by the cerebellar cortex; the results of cerebellar cortical processing are later relayed to the deep cerebellar nuclei (see Purkinje projections, below). Efferent projections from the deep cerebellar nuclei (see SCP, above) also give off collaterals that project back to the cerebellar cortex (Tolbert et al., 1978), and Purkinje cells send collaterals to nearby Purkinje cells (Palay & Chan-Palay, 2012).

#### 3.1.2. Intrinsic cerebellar pathways

The intrinsic cerebellar pathways include Purkinje corticonuclear projections, which provide the main output of the cerebellar cortex, and parallel fibers, which are the axons of granule cells and synapse onto the Purkinje cells.

##### Purkinje corticonuclear projections

###### Neuroanatomical organization

Purkinje cells are the principal output neurons of the cerebellar cortex. They form a characteristic monolayer, the Purkinje cell layer, located between the molecular layer and the granule cell layer of the cerebellar cortex. Their planar, fan-shaped dendrites extend into the molecular layer, while their axons typically descend through the granule cell layer and enter the cerebellar white matter. Purkinje cell axons terminate primarily on neurons of the deep cerebellar nuclei (dentate, emboliform, globose, and fastigial), although some also project directly to the vestibular nuclei of the brainstem. Collectively, these projections are referred to as the corticonuclear projection of the cerebellum.

A general mediolateral organization of the corticonuclear projection has been described across species. Purkinje cells in the medial vermal cortex project primarily to the fastigial nucleus, cells in the intermediate cerebellar cortex project to the globose and emboliform nuclei, and Purkinje cells in the lateral hemispheric cortex project to the dentate nucleus (Haines & Pearson, 1979; Tolbert & Bantli, 1979; Voogd et al., 1987). In addition, projections from Purkinje cells in the flocculonodular lobe directly reach the vestibular nuclei and are considered extrinsic pathways (which travel through the ICP; see Table 1). Tract-tracing studies in macaques further indicate that Purkinje cell projections are predominantly ipsilateral (Noda et al., 1990; Yamada & Noda, 1987).

The principal anatomical pathways contributing to the Purkinje corticonuclear projections are summarized in Table 5.

**Table 5.**
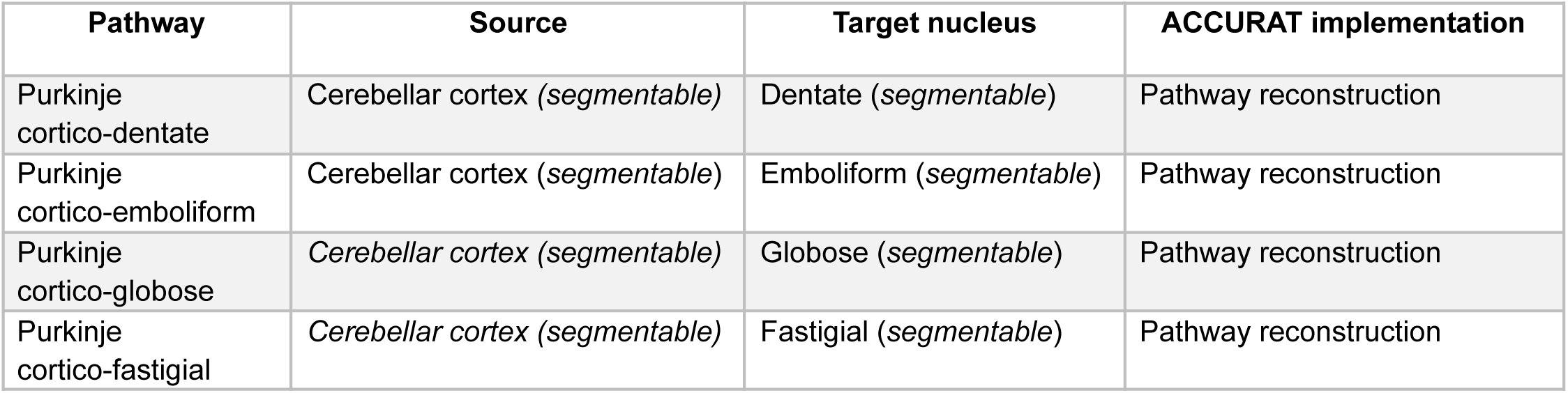
Purkinje intrinsic corticonuclear projections.

###### Tractography challenges

A principal challenge is that unconstrained tractography frequently generates streamlines that pass through, rather than terminate within, the deep cerebellar nuclei. In addition, abundant crossing fibers in cerebellar white matter (e.g., pontocerebellar projections of the MCP, ventral (anterior) spinocerebellar tract) challenge comprehensive reconstruction of Purkinje streamlines across the cerebellar cortex.

###### ACCURAT rules — Purkinje corticonuclear projections

ACCURAT identifies candidate trajectories between the cerebellar cortex and the ipsilateral deep cerebellar nuclei. Additional constraints require streamlines to remain entirely within cerebellar tissue and prevent propagation into the contralateral cerebellar hemisphere or into brainstem pathways.

##### Resulting reconstructions

In ACCURAT, these constraints recover trajectories consistent with the known anatomy of Purkinje corticonuclear projections, in which axons originate in the cerebellar cortex and terminate within the ipsilateral deep cerebellar nuclei (Figure 7).

**Figure 7.**
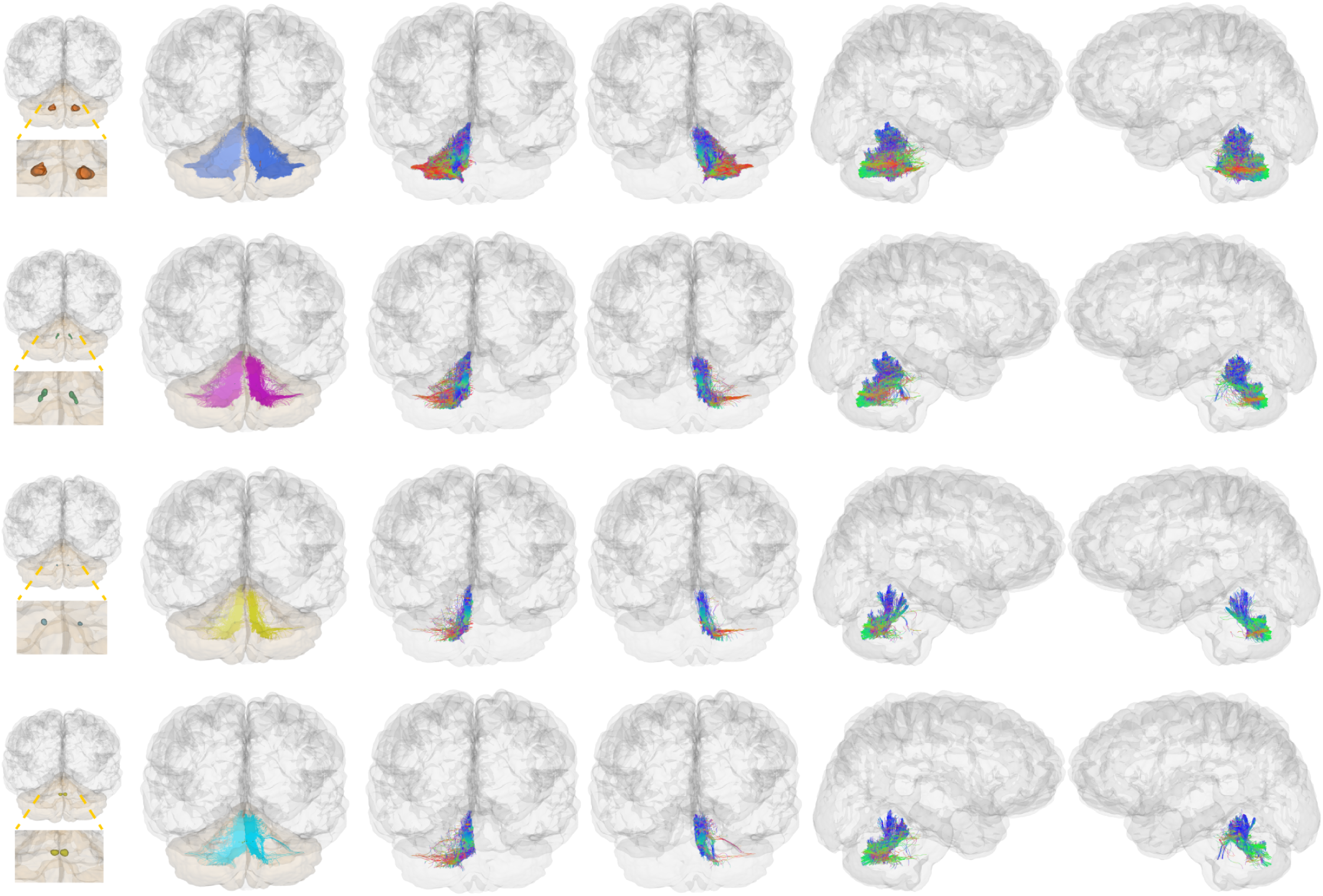
Purkinje corticonuclear projections as extracted by ACCURAT in an exemplar participant, using PTT data. Top row: Purkinje cortico-dentate; second row: Purkinje cortico-emboliform; third row: Purkinje cortico-globose; bottom row: Purkinje cortico-fastigial.

##### Intracortical parallel fibers

###### Neuroanatomical organization

Parallel fibers are the axons of granule cells. Located in the granular layer of the cerebellar cortex, the granule cells are the most abundant neurons in the mammalian brain (Herculano-Houzel, 2010). Each granule cell emits a single axon that ascends to the molecular layer of the cerebellar cortex before bifurcating into two branches that extend in opposite directions. These branches run parallel to the long axis of the cerebellar folia and are completely contained within the cerebellar cortex (Table 6).

**Table 6.** Intracortical parallel fibers.

| Pathway | Source | Target | ACCURAT implementation |
| --- | --- | --- | --- |
| Parallel fibers | Cerebellar cortex (*segmentable*) | Cerebellar cortex (*segmentable*) | Pathway reconstruction |

###### Tractography challenges

The parallel fibers have been demonstrated in ex vivo (Dell’Acqua et al., 2013; Takahashi et al., 2013) and in vivo dMRI tractography (Granziera et al., 2009; Zekelman et al., 2025). A primary challenge for tracing the parallel fibers is enabling tractography within the cerebellar cortex. This strategy differs from the cerebrum, where the cortex generally serves only as a region for initiating or terminating streamlines. Furthermore, due to partial volume effects and local propagation decisions, streamlines following the orientation of parallel fibers may exit the cerebellar cortex and enter adjacent tissues. At available image resolutions, it is not currently possible to restrict streamlines to specific layers of the cerebellar cortex (the molecular layer has a thickness of approximately 0.32 mm; (Zheng et al., 2023)). Parallel fibers have very small axonal diameters (∼0.3 µm; (Wyatt et al., 2005)) and can extend up to ∼10 mm in length (Takahashi et al., 2013), and thus tractography reconstructions can only approximate populations of parallel fibers.

###### ACCURAT rules — Parallel fibers

In the first processing step (anatomically informed cortical streamline splitting), ACCURAT performs vertex-based processing to split reconstructed streamlines at the cerebellar cortex boundary. Thus, putative parallel fiber streamlines remain entirely within the cerebellar cortex, preventing propagation into adjacent white matter or extracerebellar structures. Next, the ACCURAT query selects streamlines within the cerebellar cortex as putative reconstructions of parallel fibers.

##### Resulting reconstructions

Implemented in ACCURAT, these constraints produce trajectories consistent with the known organization of parallel fibers, which run along the cerebellar folia (Figure 8).

**Figure 8.**
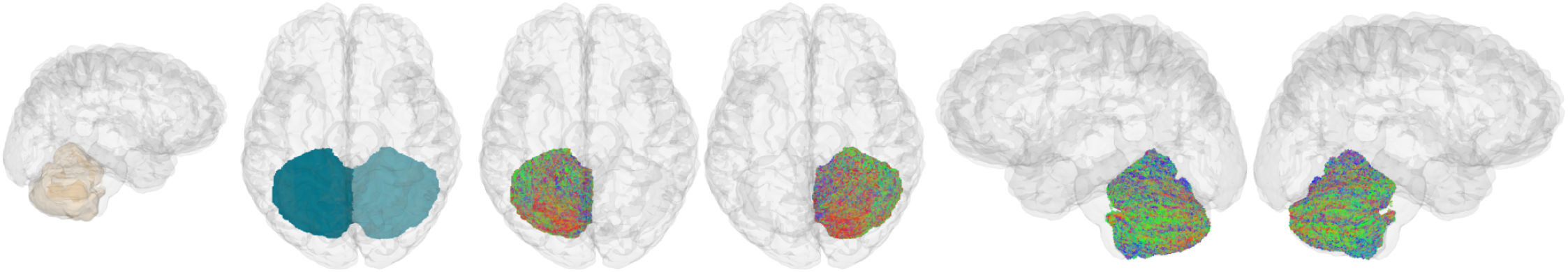
Parallel fibers as extracted by ACCURAT in an exemplar participant on PTT data.

### 3.2. Quantitative evaluation and generalization across tractography algorithms

Figure 9 shows representative reconstructions of cerebellar pathways obtained by applying ACCURAT to tractography data generated using probabilistic PTT and deterministic UKF tractography methods. Table 7 summarizes quantitative measures for each extracted pathway using PTT and UKF data, including streamline count, streamline length, and bundle identification success, measured as the number of subjects in whom at least five streamlines were identified in both hemispheres. Table S2 in the Supplementary Materials shows the bundle identification success in each hemisphere and under a stricter minimum streamline count criterion (10 instead of 5)

**Figure 9.**
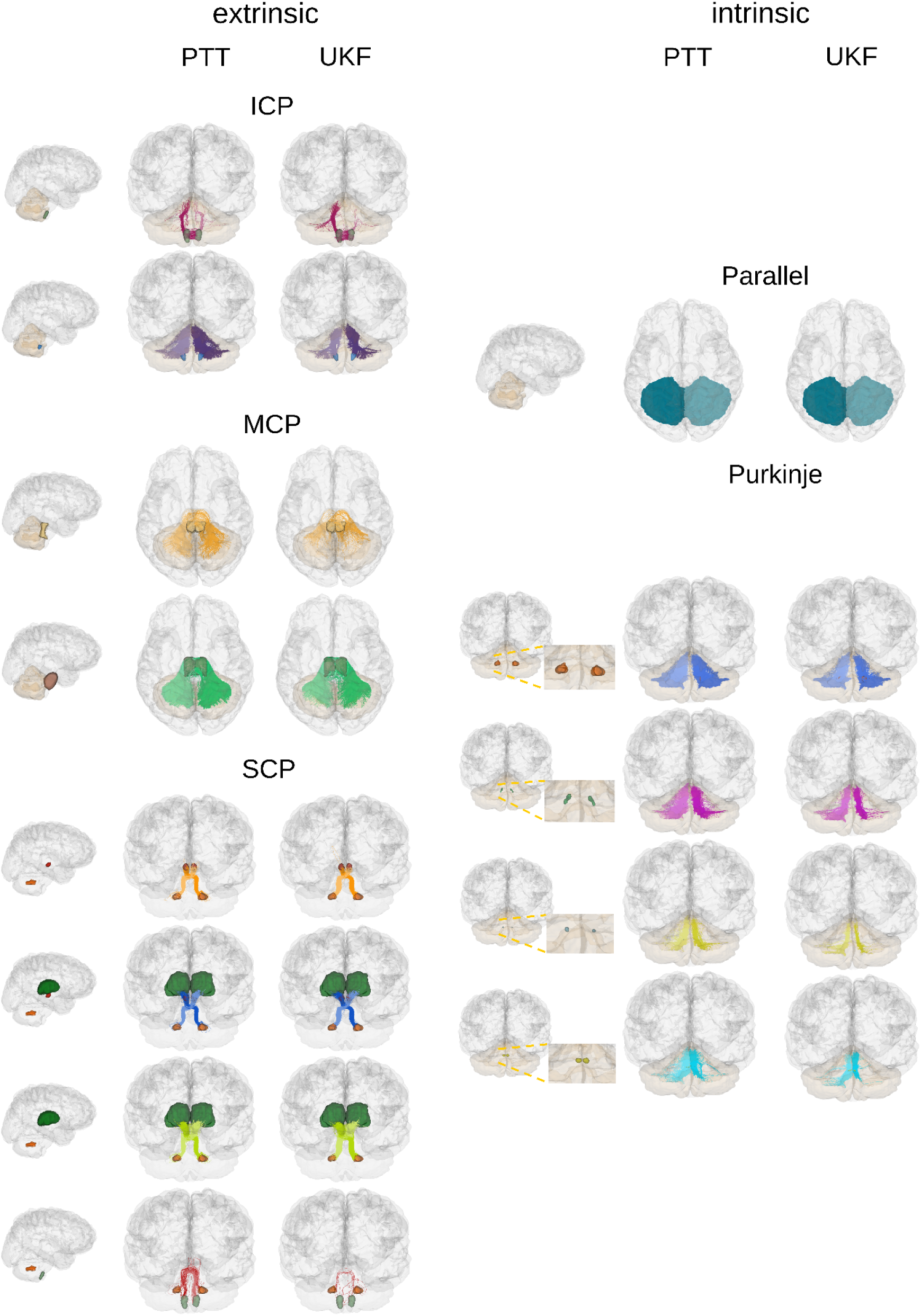
Generalization of ACCURAT across tractography algorithms. Cerebellar pathway reconstructions obtained from probabilistic parallel transport tractography (PTT) and deterministic unscented Kalman filter tractography (UKF). Despite differences in tractography algorithms, applying ACCURAT rules produces highly similar bundle reconstructions.

## 4. Discussion

Mapping cerebellar connectivity with diffusion MRI presents challenges that differ from those typically encountered in cerebral tractography, including decussations, multi-synaptic pathways, and numerous small nuclei. Furthermore, challenges known in the cerebrum, such as fiber crossings and tractography bottlenecks, are also found within the cerebellum. To begin to address these challenges, ACCURAT introduces a framework for anatomically constrained and curated reconstruction of cerebellar connectivity from diffusion MRI. By combining anatomical segmentation, densely seeded tractography, and rule-based vertex-level streamline queries, ACCURAT enables reconstruction of both extrinsic cerebellar pathways and intrinsic cerebellar circuitry. The vertex-level strategy allows anatomical constraints to be enforced directly along streamline trajectories, isolating pathway segments between nuclei while preventing propagation across synaptic boundaries. In this work, we provide both a curated set of cerebellar pathway definitions for tractography and a concise, pathway-by-pathway synthesis of cerebellar connectional anatomy based on the experimental tract-tracing literature. This synthesis component organizes anatomical evidence in a form directly useful for diffusion MRI tractography while also informing the ACCURAT rules. Using ultra-high-resolution submillimeter diffusion MRI, we demonstrate the feasibility of reconstructing multiple cerebellar pathways, including specific components of individual peduncles such as olivocerebellar projections, as well as intracerebellar intrinsic connections such as Purkinje corticonuclear pathways and parallel fibers.

### Related work in cerebellar tractography

In related work, several studies have advanced cerebellar tractography by incorporating anatomical priors, for example by selecting streamlines based on segmented nuclei (Meola, Yeh, et al., 2016) or regions of interest (Habas & Manto, 2018; Kwon et al., 2011; Nowacki et al., 2018; Palesi et al., 2015, 2017, 2020). Other studies have formed atlases of large extrinsic connections, such as the cerebellar peduncles (Tang et al., 2018; van Baarsen et al., 2016; F.-C. Yeh et al., 2018). More recent work has begun to investigate finer cerebellar pathways (Carrasco-Guerrero et al., 2025; Yin et al., 2023), subdivisions of the deep cerebellar nuclei (Steele et al., 2017; Xu et al., 2023), and topographic organization within the peduncles (Rousseau et al., 2022). Together, these studies demonstrate the value of anatomical constraints for improving the plausibility of cerebellar tractography, but they have typically focused on individual pathways or specific regions rather than providing a unified framework across intrinsic and extrinsic cerebellar pathways. For broader overviews of cerebellar tractography and cerebellar diffusion MRI, readers are referred to (Beez et al., 2021; Habas & Manto, 2018; Lundell & Steele, 2024); for more clinically focused reviews of dentato-rubro-thalamic tract targeting, see (Gravbrot et al., 2020; Lehman et al., 2020).

### ACCURAT as an open framework

ACCURAT is intended as an open framework for systematically reconstructing both intrinsic and extrinsic cerebellar connectivity and for supporting the development and evaluation of cerebellar tractography methods (see Data and Code Availability section). ACCURAT is implemented as post-processing of tractography, allowing the same anatomical rules to be applied to tractography generated using different algorithms. A practical strength of ACCURAT is that it builds on the established WMQL design of explicit query files operating on named anatomical ROIs. Because this query-based structure has already been used across multiple tractography studies, it provides a robust and extensible mechanism for defining cerebellar pathways in a transparent and reproducible way. By formalizing pathway definitions as explicit query rules, ACCURAT can support more reproducible and comparable cerebellar tractography across studies. While we have demonstrated ACCURAT’s performance on submillimeter-resolution dMRI data, larger pathways such as the cerebellar peduncles can also be reconstructed at lower resolutions, including 1.25 mm isotropic (Radwan et al., 2022) and 2 mm isotropic (Nicoletti et al., 2017).

In this study, similar reconstructions were obtained using both probabilistic PTT and deterministic UKF tractography. Detailed results of UKF tractography are presented in the Supplementary Materials, alongside additional results from a separate participant reconstructed with both PTT and UKF. The ACCURAT design facilitates comparison of tractography resulting from different methods and acquisitions and thus provides a flexible framework for systematically evaluating advances in tractography methodology (F. Zhang et al., 2025) and diffusion imaging (Lundell & Steele, 2024). Furthermore, integration of ACCURAT through automated segmentation approaches—for example, those designed for small nuclei (Legarreta et al., 2025; Olchanyi et al., 2025)—could enable automated pipelines for large-scale analyses or methodological evaluation across datasets.

### Technical challenges in cerebellar tractography

In the Results, we described pathway-specific tractography challenges with reference to the literature; here, we discuss additional challenges that emerged during ACCURAT development.

One challenge arises from defining rules for the reconstruction of bidirectional connections between nuclei, where two different pathways may share the same endpoints. For example, the deep cerebellar nuclei project via the SCP to the contralateral inferior olive (dentato-olivary pathway), while the inferior olive sends projections via the contralateral ICP to the cerebellar cortex, with collateral branches to the deep cerebellar nuclei (olivo-cerebellar pathway). These two pathways are not separable based on endpoint criteria alone, because both have streamline endpoints in the dentate and inferior olive. Therefore, additional approaches such as fiber clustering or “waypoint”-style inclusion ROIs are needed to enable more specific tracing of particular cerebellar circuits. In ACCURAT, we addressed this by using an additional ROI to extract the dentato-olivary pathway, excluding streamlines that passed through the ipsilateral ICP ROI. This endpoint non-specificity is a characteristic of multiple cerebellar circuits and is also a challenge in developing connectivity matrices for research into the cerebellar connectome, as traditionally such matrices only consider streamline endpoints.

A further challenge in developing rules for cerebellar tractography is that some nuclei involved in cerebellar pathways cannot be reliably visualized, even in submillimeter diffusion MRI, due to their small size and limited MRI contrast. For example, pontocerebellar projections originate in numerous pontine nuclei that are not individually visible in MRI. In ACCURAT, this issue was addressed using regions of the pons as bounding anatomical regions, enabling the development of queries to define two components of the MCP. Future work could apply this strategy to investigate additional pathways beyond the scope of the present study. Another possible strategy would be fiber clustering to separate putative components within larger pathways. Multimodal imaging may also improve visualization and segmentation of small nuclei, although inter-modality registration remains challenging for small brainstem structures.

We also observed systematic biases and practical reconstruction challenges in cerebellar tractography. The Purkinje fibers are anatomically expected to originate in the entire cerebellar cortex and project to the deep cerebellar nuclei; however, we observed that tracing of streamlines from lateral cortical regions was sparse, making cerebellar cortical coverage non-uniform. These Purkinje streamlines would need to cross other cerebellar pathways to reach the dentate nucleus. Thus, the observed coverage bias was likely related to challenges in depicting crossing and/or kissing fibers, such as the MCP that traverses this region. In addition to the well-known challenges in tracing the SCP decussation (Coenen et al., 2021), we observed that other less-studied decussating pathways are also challenging to trace. These included the descending dentato-olivary pathway of the SCP and the olivocerebellar pathway of the ICP, which were the smallest bundles included in ACCURAT (by number of streamlines; see Table 7). On these challenging bundles, PTT was generally more robust at tracing the dentato-olivary SCP, while both tractography methods struggled to reconstruct the olivocerebellar pathway.

**Table 7.** Quantitative reconstruction of cerebellar pathways across tractography algorithms. Mean ± standard deviation of streamline counts and streamline lengths for each reconstructed bundle using probabilistic parallel transport tractography (PTT) and deterministic unscented Kalman filter tractography (UKF). The number of subjects for which each bundle was successfully reconstructed (with 5 or more streamlines in both hemispheres) is also reported.

| Pathway | PTT |  |  | UKF |  |  |
| --- | --- | --- | --- | --- | --- | --- |
|  | Streamline count | Length (mm) | Subjects | Streamline count | Length (mm) | Subjects |
| ***Extrinsic cerebellar pathways*** |  |  |  |  |  |  |
| **ICP olivocerebellar** | 63 ± 108 | 61.48 ± 12.07 | 6 | 234 ± 203 | 72.05 ± 15.57 | 8 |
| **ICP peduncle** | 10157 ± 2276 | 75.49 ± 5.04 | 9 | 6393 ± 1636 | 58.54 ± 3.5 | 9 |
| **MCP pontocerebellar** | 28199 ± 6304 | 112.63 ± 3.84 | 9 | 7091 ± 1066 | 96.11 ± 3.83 | 9 |
| **MCP reticulocerebellar** | 2025 ± 943 | 102.35 ± 7.21 | 9 | 896 ± 265 | 73.84 ± 6.08 | 9 |
| **SCP dentato-rubral** | 1848 ± 1014 | 86.63 ± 4.49 | 9 | 6710 ± 2374 | 82.2 ± 3.16 | 9 |
| **SCP dentato-rubro-thalamic** | 796 ± 569 | 110.95 ± 8.0 | 9 | 4372 ± 1503 | 103.96 ± 4.09 | 9 |
| **SCP dentato-thalamic** | 1051 ± 910 | 112.34 ± 7.38 | 9 | 4504 ± 1980 | 103.71 ± 3.66 | 9 |
| **SCP dentato-olivary** | 380 ± 460 | 141.41 ± 6.86 | 9 | 60 ± 31 | 114.8 ± 20.72 | 9 |
| ***Intrinsic cerebellar pathways*** |  |  |  |  |  |  |
| **Purkinje cortico-dentate** | 47800 ± 12050 | 27.09 ± 1.93 | 9 | 40426 ± 6943 | 18.53 ± 1.79 | 9 |
| **Purkinje cortico-emboliform** | 11176 ± 2739 | 31.09 ± 2.13 | 9 | 10030 ± 2692 | 25.86 ± 1.96 | 9 |
| **Purkinje cortico-globose** | 8191 ± 2185 | 25.84 ± 3.43 | 9 | 6231 ± 1653 | 19.59 ± 3.51 | 9 |
| **Purkinje cortico-fastigial** | 10665 ± 2062 | 19.31 ± 2.54 | 9 | 6044 ± 1608 | 17.04 ± 3.82 | 9 |
| **Parallel fibers** | 1254271 ± 67673 | 51.17 ± 3.39 | 9 | 897697 ± 62823 | 40.63 ± 2.66 | 9 |

An additional challenge arises from collateral branches, which remain poorly characterized in the macaque and human cerebellum. Relatively little is known about whether they are consistently present at particular locations or how they vary in length, branching pattern, or incidence. Given this complexity, we did not attempt to trace collateral branches in this study. We did, however, observe some olivo-dentate streamlines that were consistent with the anatomy of established collateral branches of olivocerebellar axons traveling through the ICP. We removed these to isolate the dentato-olivary pathway, but future work should investigate whether relatively large and consistent collateral connections can be reconstructed using dMRI, as suggested by a recent 7T reconstruction of the olivo-dentate ICP pathway in a patient (Calzoni et al., 2025).

### Challenges in neuroanatomical ground truth for cerebellar pathways

We have provided an overview of known cerebellar pathways to guide tractography development, including the ACCURAT framework proposed here. However, the corpus of neuroanatomical knowledge about the primate cerebellum is surprisingly sparse (see (Voogd, 2003)). Most of what is known about cerebellar connectivity was assembled by Larsell, Jansen, Brodal, and others largely between the 1920s and the early 1970s, predominantly in the cat, using degeneration approaches, in which experimental lesions induced Wallerian or retrograde degeneration to reveal afferent and efferent connections (Jansen & Brodal, 1954; Larsell & Jansen, 1972). These investigators mapped cerebellar circuits in considerable detail, but the degeneration technique had important limitations, including poor visualization of small axons and collateral branches, and confounds from vascular damage, fibers of passage, and collateral injury. As a result, the degeneration technique was largely abandoned once anterograde and retrograde tracer methods became available in the early 1970s.

Modern experimental tract-tracing methods increased sensitivity, specificity, and precision while eliminating the principal confounds of degeneration. Despite these advances, cerebellar circuits received far less attention than forebrain circuits, and primate tract-tracing cerebellar work in particular has remained limited. Only approximately 50 tract-tracing studies have been performed on the macaque cerebellum, most focused on corticopontine projections or cerebello-thalamic connections. Consequently, the organization of cerebellar circuits in the primate brain remains poorly resolved, with current models relying on extrapolation from non-primate species. Such extrapolation is likely reasonable for phylogenetically older regions of the cerebellum, such as the vermis and flocculonodular lobe, but becomes increasingly problematic in regions that have expanded in primates (Sereno et al., 2020), particularly the lateral hemispheres and crus I and II, where direct macaque data remain limited. These gaps in neuroanatomical ground truth present an additional challenge for cerebellar tractography, because both pathway definitions and the interpretation of reconstructed bundles depend on anatomical evidence that is incomplete for many primate cerebellar circuits.

### Limitations and future work

In this work, we focused on intrinsic and extrinsic cerebellar pathways; however, cerebellar pathways participate in distributed circuits that extend well beyond the cerebellum itself. The ACCURAT framework could be straightforwardly extended to reconstruct pathways beyond the cerebellum, such as pathways that synapse on pontine nuclei (e.g., cortico-pontine, hypothalamo-pontine) and rubro-olivary pathways. Cerebellar pathways not implemented here generally involved practical challenges of anatomical delineation, small size, or complex trajectories rather than conceptual limitations of the framework itself. Thus, we provided candidate definitions and a concise synthesis of the experimental tract tracing literature to support future work studying additional pathways. The ACCURAT system can also be extended with finer pathway definitions, such as lobule-specific cerebellar cortex connectivity, which can be readily implemented within the framework in the future. Even at submillimeter resolution, we find that reconstruction of some bundles remains challenging, highlighting both the promise of high-resolution diffusion imaging and the limitations of current approaches. Future work is needed to determine how improvements in spatial resolution, tractography methods, fiber modeling, and anatomical constraints can further improve the reconstruction of these pathways.

## 5. Conclusion

In this work, we introduced ACCURAT, an open framework for anatomically constrained and curated reconstruction of cerebellar connectivity from diffusion MRI. Using ultra-high-resolution submillimeter diffusion MRI and vertex-level anatomical rules, we demonstrate the feasibility of reconstructing multiple intrinsic and extrinsic cerebellar pathways in vivo. In parallel, we provide a pathway-by-pathway neuroanatomical synthesis that consolidates dispersed experimental evidence into a tractography-oriented resource for the field. Together, the framework and synthesis establish a practical foundation for anatomically informed cerebellar tractography and for future investigation of cerebellar circuits in health and disease.

## Data and Code Availability

The ACCURAT query-based streamline processing framework will be made available as open source at https://github.com/SlicerDMRI/accurat. MRI data will be shared via the National Institute of Mental Health Data Archive (https://nda.nih.gov/edit_collection.html?id=3950).

## Author Contributions

**Jon Haitz Legarreta**: Conceptualization, Methodology, Data Curation, Investigation, Software, Formal analysis, Validation, Writing - Original Draft, Writing - Review & Editing, Visualization, Funding acquisition; **Richard J. Rushmore**: Conceptualization, Methodology, Data Curation, Investigation, Validation, Writing - Original Draft, Writing - Review & Editing, Funding acquisition; **Edward H. Yeterian**: Methodology, Validation, Writing - Review & Editing; **Nikos Makris**: Methodology, Validation, Writing - Review & Editing, Funding acquisition; **Yogesh Rathi**: Conceptualization, Methodology, Investigation, Resources, Writing - Review & Editing, Funding acquisition; **Lauren J. O’Donnell**: Conceptualization, Methodology, Data Curation, Validation, Writing - Original Draft, Writing - Review & Editing, Supervision, Project administration, Funding acquisition.

## Funding

Research supported by the National Institute of Mental Health and National Institute of Neurological Disorders and Stroke of the National Institute of Health under Award Numbers R01MH132610, R01MH125860, R01MH119222, R01NS125307, R01NS125781, R01MH116173, R01MH112748, R21NS136960. The content is solely the responsibility of the authors and does not necessarily represent the official views of the National Institute of Health.

## Declaration of Competing Interests

Authors declare no competing interests.

## Acknowledgements

We thank the Enterprise Research Infrastructure & Services at Mass General Brigham for their support and for providing the Scientific Computing (SciC) Linux Clusters. This work used the Jetstream2 on-demand computing infrastructure (Hancock et al., 2021) at Indiana University through allocation MED230035 from the Advanced Cyberinfrastructure Coordination Ecosystem: Services & Support (ACCESS) program, which is supported by National Science Foundation grants #2138259, #2138286, #2138307, #2137603, and #2138296 (Boerner et al., 2023).

## Notes

### Competing Interest Statement

The authors have declared no competing interest.

## References

Amino, Y., Kyuhou, S., Matsuzaki, R., & Gemba, H. (2001). Cerebello-thalamo-cortical projections to the posterior parietal cortex in the macaque monkey. Neuroscience Letters, 309(1), 29–32.

Asanuma, C., Thach, W. R., & Jones, E. G. (1983a). Anatomical evidence for segregated focal groupings of efferent cells and their terminal ramifications in the cerebellothalamic pathway of the monkey. Brain Research, 286(3), 267–297.

Asanuma, C., Thach, W. T., & Jones, E. G. (1983b). Brainstem and spinal projections of the deep cerebellar nuclei in the monkey, with observations on the brainstem projections of the dorsal column nuclei. Brain Research, 286(3), 299–322.

Asanuma, C., Thach, W. T., & Jones, E. G. (1983c). Distribution of cerebellar terminations and their relation to other afferent terminations in the ventral lateral thalamic region of the monkey. Brain Research, 286(3), 237–265.

Aydogan, D. B., Jacobs, R., Dulawa, S., Thompson, S. L., Francois, M. C., Toga, A. W., Dong, H., Knowles, J. A., & Shi, Y. (2018). When tractography meets tracer injections: a systematic study of trends and variation sources of diffusion-based connectivity. Brain Structure & Function, 223(6), 2841–2858.

Aydogan, D. B., & Shi, Y. (2021). Parallel transport tractography. IEEE Transactions on Medical Imaging, 40(2), 635–647.

Balsters, J. H., Cussans, E., Diedrichsen, J., Phillips, K. A., Preuss, T. M., Rilling, J. K., & Ramnani, N. (2010). Evolution of the cerebellar cortex: the selective expansion of prefrontal-projecting cerebellar lobules. NeuroImage, 49(3), 2045–2052.

Barati Shoorche, A., Farnia, P., Makkiabadi, B., & Leemans, A. (2025). A review on learning-based algorithms for tractography and human brain white matter tracts recognition. Neuroradiology, 67(8), 2041–2067.

Batton, R. R., 3rd, Jayaraman, A., Ruggiero, D., & Carpenter, M. B. (1977). Fastigial efferent projections in the monkey: an autoradiographic study. The Journal of Comparative Neurology, 174(2), 281–305.

Beez, T., Munoz-Bendix, C., Steiger, H.-J., & Hänggi, D. (2021). Functional tracts of the cerebellum-essentials for the neurosurgeon. Neurosurgical Review, 44(1), 273–278.

Boerner, T. J., Deems, S., Furlani, T. R., Knuth, S. L., & Towns, J. (2023). ACCESS: Advancing Innovation: NSF’s Advanced Cyberinfrastructure Coordination Ecosystem: Services & Support. Practice and Experience in Advanced Research Computing, 173–176.

Bostan, A. C., & Strick, P. L. (2013). Cerebellar Outputs in Non-human Primates: An Anatomical Perspective Using Transsynaptic Tracers. In Handbook of the Cerebellum and Cerebellar Disorders (pp. 549–569). 10.1007/978-94-007-1333-8_25

Brodal, A., & Brodal, P. (1985). Observations on the secondary vestibulocerebellar projections in the macaque monkey. Experimental Brain Research, 58(1), 62–74.

Brodal, P. (1979). The pontocerebellar projection in the rhesus monkey: an experimental study with retrograde axonal transport of horseradish peroxidase. Neuroscience, 4(2), 193–208.

Brodal, P. (1980). The projection from the nucleus reticularis tegmenti pontis to the cerebellum in the rhesus monkey. Experimental Brain Research, 38(1), 29–36.

Brodal, P. (1982). Further observations on the cerebellar projections from the pontine nuclei and the nucleus reticularis tegmenti pontis in the rhesus monkey. The Journal of Comparative Neurology, 204(1), 44–55.

Brodal, P., & Brodal, A. (1981). The olivocerebellar projection in the monkey. Experimental studies with the method of retrograde tracing of horseradish peroxidase. The Journal of Comparative Neurology, 201(3), 375–393.

Brogna, C., Perera, N., Ghimire, P., Bruchhage, M. M. K., Abela, E., Richardson, M. P., Vergani, F., Bhangoo, R., & Ashkan, K. (2022). First human in vivo neuroelectrophysiology recordings of uncrossed dentatothalamocortical white-matter connections: On the fast tract: On the fast tract. Neurology, 99(8), 332–335.

Buckner, R. L. (2013). The cerebellum and cognitive function: 25 years of insight from anatomy and neuroimaging. Neuron, 80(3), 807–815.

Calzoni, T., Donatelli, G., Migaleddu, G., Lancione, M., Cecchi, P., Biagi, L., Caniglia, M., Ceravolo, R., & Cosottini, M. (2025). Hypertrophic olivary degeneration: a 7 Tesla advanced imaging case report. Frontiers in Neuroscience, 19(1656655), 1656655.

Carpenter, M. B., & Stevens, G. H. (1957). Structural and functional relationships between the deep cerebellar nuclei and the brachium conjunctivum in the rhesus monkey. The Journal of Comparative Neurology, 107(1), 109–163.

Carpenter, M. B., & Sutin, J. (1983). Human neuroanatomy, 8th *ed.* Williams & Wilkins.

Carrasco-Guerrero, D., Riquelme, P., Hernández, C., & Guevara, P. (2025). Atlas of Cerebellar Connections Based on ROIs and dMRI Tractography. 2025 21st International Symposium on Biomedical Image Processing and Analysis (SIPAIM), 1–4.

Çavdar, S., Esen Aydın, A., Algin, O., & Aydoğmuş, E. (2020). Fiber dissection and 3-tesla diffusion tensor tractography of the superior cerebellar peduncle in the human brain: emphasize on the cerebello-hypthalamic fibers. Brain Structure & Function, 225(1), 121–128.

Chan-Palay, V. (1977). The cerebellar dentate nucleus. In Cerebellar Dentate Nucleus (pp. 1–24). Springer Berlin Heidelberg.

Coenen, V. A., Sajonz, B. E., Reinacher, P. C., Kaller, C. P., Urbach, H., & Reisert, M. (2021). A detailed analysis of anatomical plausibility of crossed and uncrossed streamline rendition of the dentato-rubro-thalamic tract (DRT(T)) in a commercial stereotactic planning system. Acta Neurochirurgica, 163(10), 2809–2824.

Coudé, D., Parent, A., & Parent, M. (2018). Single-axon tracing of the corticosubthalamic hyperdirect pathway in primates. Brain Structure & Function, 223(9), 3959–3973.

Dell’Acqua, F., Bodi, I., Slater, D., Catani, M., & Modo, M. (2013). MR Diffusion Histology and Micro-Tractography Reveal Mesoscale Features of the Human Cerebellum. Cerebellum, 12(6), 923–931.

Fatemi, S. H., Aldinger, K. A., Ashwood, P., Bauman, M. L., Blaha, C. D., Blatt, G. J., Chauhan, A., Chauhan, V., Dager, S. R., Dickson, P. E., Estes, A. M., Goldowitz, D., Heck, D. H., Kemper, T. L., King, B. H., Martin, L. A., Millen, K. J., Mittleman, G., Mosconi, M. W., … Welsh, J. P. (2012). Consensus paper: pathological role of the cerebellum in autism. Cerebellum, 11(3), 777–807.

Fedorov, A., Beichel, R., Kalpathy-Cramer, J., Finet, J., Fillion-Robin, J.-C., Pujol, S., Bauer, C., Jennings, D., Fennessy, F., Sonka, M., Buatti, J., Aylward, S., Miller, J. V., Pieper, S., & Kikinis, R. (2012). 3D Slicer as an image computing platform for the Quantitative Imaging Network. Magnetic Resonance Imaging, 30(9), 1323–1341.

Fischl, B. (2012). FreeSurfer. NeuroImage, 62(2), 774–781.

Gaviraghi, M., Savini, G., Castellazzi, G., Palesi, F., Rolandi, N., Sacco, S., Pichiecchio, A., Mariani, V., Tartara, E., Tassi, L., Vitali, P., D’Angelo, E., & Wheeler-Kingshott, C. A. M. G. (2021). Automatic Segmentation of Dentate Nuclei for Microstructure Assessment: Example of Application to Temporal Lobe Epilepsy Patients. Computational Diffusion MRI, 263–278.

Gellersen, H. M., Guell, X., & Sami, S. (2021). Differential vulnerability of the cerebellum in healthy ageing and Alzheimer’s disease. NeuroImage. Clinical, 30, 102605.

Gonzalo-Ruiz, A., Leichnetz, G. R., & Smith, D. J. (1988). Origin of cerebellar projections to the region of the oculomotor complex, medial pontine reticular formation, and superior colliculus in new world monkeys: A retrograde horseradish peroxidase study. In The Journal of Comparative Neurology (Vol. 268, Issue 4, pp. 508–526). 10.1002/cne.902680404

Granziera, C., Schmahmann, J. D., Hadjikhani, N., Meyer, H., Meuli, R., Wedeen, V., & Krueger, G. (2009). Diffusion spectrum imaging shows the structural basis of functional cerebellar circuits in the human cerebellum in vivo. PloS One, 4(4), e5101.

Gravbrot, N., Saranathan, M., Pouratian, N., & Kasoff, W. S. (2020). Advanced imaging and direct targeting of the motor thalamus and dentato-rubro-thalamic tract for tremor: A systematic review. Stereotactic and Functional Neurosurgery, 98(4), 220–240.

Groenewegen, H. J., Voogd, J., & Freedman, S. L. (1979). The parasagittal zonation within the olivocerebellar projection. II. Climbing fiber distribution in the intermediate and hemispheric parts of cat cerebellum. The Journal of Comparative Neurology, 183(3), 551–601.

Habas, C., & Manto, M. (2018). Probing the neuroanatomy of the cerebellum using tractography. Handbook of Clinical Neurology, 154, 235–249.

Haines, D. E., May, P. J., & Dietrichs, E. (1990). Neuronal connections between the cerebellar nuclei and hypothalamus in Macaca fascicularis: cerebello-visceral circuits. The Journal of Comparative Neurology, 299(1), 106–122.

Haines, D. E., & Pearson, J. C. (1979). Cerebellar corticonuclear - nucleocortical topography: a study of the tree shrew (Tupaia) paraflocculus. The Journal of Comparative Neurology, 187(4), 745–758.

Hancock, D. Y., Fischer, J., Lowe, J. M., Snapp-Childs, W., Pierce, M., Marru, S., Coulter, J. E., Vaughn, M., Beck, B., Merchant, N., Skidmore, E., & Jacobs, G. (2021). Jetstream2: Accelerating cloud computing via Jetstream. *Practice and Experience in Advanced Research Computing*, Article Article 11.

Harrison, O. K., Guell, X., Klein-Flügge, M. C., & Barry, R. L. (2021). Structural and resting state functional connectivity beyond the cortex. NeuroImage, 240, 118379.

Herculano-Houzel, S. (2010). Coordinated scaling of cortical and cerebellar numbers of neurons. Frontiers in Neuroanatomy, 4, 12.

Hoffmann, M., Billot, B., Greve, D. N., Iglesias, J. E., Fischl, B., & Dalca, A. V. (2022). SynthMorph: Learning contrast-invariant registration without acquired images. IEEE Transactions on Medical Imaging, 41(3), 543–558.

Hoffmann, M., Hoopes, A., Greve, D. N., Fischl, B., & Dalca, A. V. (2024). Anatomy-aware and acquisition-agnostic joint registration with SynthMorph. Imaging Neuroscience (Cambridge, Mass.), 2, 1–33.

Hoshi, E., Tremblay, L., Féger, J., Carras, P. L., & Strick, P. L. (2005). The cerebellum communicates with the basal ganglia. In Nature Neuroscience (Vol. 8, Issue 11, pp. 1491–1493). 10.1038/nn1544

Ikeda, Y., Noda, H., & Sugita, S. (1989). Olivocerebellar and cerebelloolivary connections of the oculomotor region of the fastigial nucleus in the macaque monkey. The Journal of Comparative Neurology, 284(3), 463–488.

Ilinsky, I. A., & Kultas-Ilinsky, K. (1984). An autoradiographic study of topographical relationships between pallidal and cerebellar projections to the cat thalamus. Experimental Brain Research, 54(1), 95–106.

Ilinsky, I. A., Kultas-Ilinsky, K., Rosina, A., & Haddy, M. (1987). Quantitative evaluation of crossed and uncrossed projections from basal ganglia and cerebellum to the cat thalamus. Neuroscience, 21(1), 207–227.

Jacobs, H. I. L., Hopkins, D. A., Mayrhofer, H. C., Bruner, E., van Leeuwen, F. W., Raaijmakers, W., & Schmahmann, J. D. (2018). The cerebellum in Alzheimer’s disease: evaluating its role in cognitive decline. In Brain (Vol. 141, Issue 1, pp. 37–47). 10.1093/brain/awx194

Jansen, J., & Brodal, A. (1954). Aspects of cerebellar anatomy. Edited by Jan Jansen and Alf Brodal, Oslo, Johan Grundt Tanum. 1954. 423 p. Price $9.00. *(Reviewed by* Gerhardt von Bonin). The Journal of Comparative Neurology, 101(3), 830–834.

Jenkinson, M., Beckmann, C. F., Behrens, T. E. J., Woolrich, M. W., & Smith, S. M. (2012). FSL. NeuroImage, 62(2), 782–790.

Jeong, J.-W., Tiwari, V. N., Behen, M. E., Chugani, H. T., & Chugani, D. C. (2014). In vivo detection of reduced Purkinje cell fibers with diffusion MRI tractography in children with autistic spectrum disorders. Frontiers in Human Neuroscience, 8, 110.

Jeurissen, B., Leemans, A., Tournier, J.-D., Jones, D. K., & Sijbers, J. (2013). Investigating the prevalence of complex fiber configurations in white matter tissue with diffusion magnetic resonance imaging. Human Brain Mapping, 34(11), 2747–2766.

Kalil, K. (1979). Projections of the cerebellar and dorsal column nuclei upon the inferior olive in the rhesus monkey: an autoradiographic study. The Journal of Comparative Neurology, 188(1), 43–62.

Kim, J., Patriat, R., Kaplan, J., Solomon, O., & Harel, N. (2020). Deep Cerebellar Nuclei Segmentation via Semi-Supervised Deep Context-Aware Learning from 7T Diffusion MRI. *IEEE Access : Practical Innovations*, Open Solutions, 8, 101550–101568.

Kita, T., & Kita, H. (2012). The subthalamic nucleus is one of multiple innervation sites for long-range corticofugal axons: A single-axon tracing study in the rat. The Journal of Neuroscience: The Official Journal of the Society for Neuroscience, 32, 5990–5999.

Kwon, H. G., Hong, J. H., Hong, C. P., Lee, D. H., Ahn, S. H., & Jang, S. H. (2011). Dentatorubrothalamic tract in human brain: diffusion tensor tractography study. Neuroradiology, 53(10), 787–791.

Kyuhou, S., Matsuzaki, R., & Gemba, H. (1997). Cerebello-cerebral projections onto the ventral part of the frontal cortex of the macaque monkey. Neuroscience Letters, 230(2), 101–104.

Langer, T., Fuchs, A. F., Scudder, C. A., & Chubb, M. C. (1985). Afferents to the flocculus of the cerebellum in the rhesus macaque as revealed by retrograde transport of horseradish peroxidase. The Journal of Comparative Neurology, 235(1), 1–25.

Larsell, O., & Jansen, J. (1972). The Comparative Anatomy and Histology of the Cerebellum. University of Minnesota Press.

Legarreta, J. H., Lan, Z., Chen, Y., Zhang, F., Yeterian, E. H., Makris, N., Rushmore, R. J., Rathi, Y., & O’Donnell, L. J. (2025). Towards an informed choice of diffusion MRI image contrasts for cerebellar segmentation. Human Brain Mapping, 46(11), e70317.

Lehman, V. T., Lee, K. H., Klassen, B. T., Blezek, D. J., Goyal, A., Shah, B. R., Gorny, K. R., Huston, J., & Kaufmann, T. J. (2020). MRI and tractography techniques to localize the ventral intermediate nucleus and dentatorubrothalamic tract for deep brain stimulation and MR-guided focused ultrasound: a narrative review and update. Neurosurgical Focus, 49(1), E8.

Leitner, Y., Travis, K. E., Ben-Shachar, M., Yeom, K. W., & Feldman, H. M. (2015). Tract profiles of the cerebellar white matter pathways in children and adolescents. *Cerebellum (London*, England), 14(6), 613–623.

Lundell, H., & Steele, C. J. (2024). Cerebellar imaging with diffusion magnetic resonance imaging: approaches, challenges, and potential. Current Opinion in Behavioral Sciences, 56(101353), 101353.

Luo, Y., Fujita, H., Nedelescu, H., Biswas, M. S., Sato, C., Ying, S., Takahashi, M., Akita, K., Higashi, T., Aoki, I., & Sugihara, I. (2017). Lobular homology in cerebellar hemispheres of humans, non-human primates and rodents: a structural, axonal tracing and molecular expression analysis. Brain Structure & Function, 222(6), 2449–2472.

MacLeod, C. (2012). The missing link: evolution of the primate cerebellum. Progress in Brain Research, 195, 165–187.

Magielse, N., Toro, R., Steigauf, V., Abbaspour, M., Eickhoff, S. B., Heuer, K., & Valk, S. L. (2023). Phylogenetic comparative analysis of the cerebello-cerebral system in 34 species highlights primate-general expansion of cerebellar crura I-II. Communications Biology, 6(1), 1188.

Maier-Hein, K. H., Neher, P. F., Houde, J.-C., Côté, M.-A., Garyfallidis, E., Zhong, J., Chamberland, M., Yeh, F.-C., Lin, Y.-C., Ji, Q., Reddick, W. E., Glass, J. O., Chen, D. Q., Feng, Y., Gao, C., Wu, Y., Ma, J., He, R., Li, Q., … Descoteaux, M. (2017). The challenge of mapping the human connectome based on diffusion tractography. Nature Communications, 8(1), 1349.

Makris, N., Hodge, S. M., Haselgrove, C., Kennedy, D. N., Dale, A., Fischl, B., Rosen, B. R., Harris, G., Caviness, V. S., Jr, & Schmahmann, J. D. (2003). Human cerebellum: surface-assisted cortical parcellation and volumetry with magnetic resonance imaging. Journal of Cognitive Neuroscience, 15(4), 584–599.

Makris, N., Schlerf, J. E., Hodge, S. M., Haselgrove, C., Albaugh, M. D., Seidman, L. J., Rauch, S. L., Harris, G., Biederman, J., Caviness, V. S., Jr, Kennedy, D. N., & Schmahmann, J. D. (2005). MRI-based surface-assisted parcellation of human cerebellar cortex: an anatomically specified method with estimate of reliability. NeuroImage, 25(4), 1146–1160.

Martinez, J. D., Mooshagian, E., Wang, H., Karlsgodt, K. H., Uddin, L. Q., & Nomi, J. S. (2026). Diffusion tensor imaging of cerebellar peduncles and association tracts in a case of primary complete callosal agenesis. Brain Structure & Function, 231(3). 10.1007/s00429-026-03091-y

Mason, A., Ilinsky, I. A., Maldonado, S., & Kultas-Ilinsky, K. (2000). Thalamic terminal fields of individual axons from the ventral part of the dentate nucleus of the cerebellum in Macaca mulatta. The Journal of Comparative Neurology, 421(3), 412–428.

Matsushita, M., & Ikeda, M. (1970). Olivary projections to the cerebellar nuclei in the cat. Experimental Brain Research, 10(5), 488–500.

May, P. J., Hartwich-Young, R., Nelson, J., Sparks, D. L., & Porter, J. D. (1990). Cerebellotectal pathways in the macaque: implications for collicular generation of saccades. Neuroscience, 36(2), 305–324.

May, P. J., Porter, J. D., & Gamlin, P. D. (1992). Interconnections between the primate cerebellum and midbrain near-response regions. The Journal of Comparative Neurology, 315(1), 98–116.

Meola, A., Comert, A., Yeh, F. C., Sivakanthan, S., & Fernandez-Miranda, J. C. (2016). The nondecussating pathway of the dentatorubrothalamic tract in humans: human connectome-based tractographic study and microdissection validation. Journal of Neurosurgery, 124(5), 1406–1412.

Meola, A., Yeh, F.-C., Fellows-Mayle, W., Weed, J., & Fernandez-Miranda, J. C. (2016). Human Connectome-Based Tractographic Atlas of the Brainstem Connections and Surgical Approaches. Neurosurgery, 79(3), 437–455.

Nagao, S., Kitamura, T., Nakamura, N., Hiramatsu, T., & Yamada, J. (1997a). Differences of the primate flocculus and ventral paraflocculus in the mossy and climbing fiber input organization. The Journal of Comparative Neurology, 382(4), 480–498.

Nagao, S., Kitamura, T., Nakamura, N., Hiramatsu, T., & Yamada, J. (1997b). Location of efferent terminals of the primate flocculus and ventral paraflocculus revealed by anterograde axonal transport methods. Neuroscience Research, 27(3), 257–269.

Nakano, K., Takimoto, T., Kayahara, T., Takeuchi, Y., & Kobayashi, Y. (1980). Distribution of cerebellothalamic neurons projecting to the ventral nuclei of the thalamus: an HRP study in the cat. The Journal of Comparative Neurology, 194(2), 427–439.

Nicoletti, G., Caligiuri, M. E., Cherubini, A., Morelli, M., Novellino, F., Arabia, G., Salsone, M., & Quattrone, A. (2017). A fully automated, atlas-based approach for superior cerebellar peduncle evaluation in progressive supranuclear palsy phenotypes. AJNR. American Journal of Neuroradiology, 38(3), 523–530.

Nieuwenhuys, R., Voogd, J., & van Huijzen, C. (2008). The Human Central Nervous System. Springer.

Noda, H., Sugita, S., & Ikeda, Y. (1990). Afferent and efferent connections of the oculomotor region of the fastigial nucleus in the macaque monkey. The Journal of Comparative Neurology, 302(2), 330–348.

Noguera, C. B., Bao, S., Petersen, K. J., Lopez, A. M., Reid, J. A., Plassard, A. J., Zald, D. H., Claassen, D. O., Dawant, B. M., & Landman, B. A. (2019). Using deep learning for a diffusion-based segmentation of the dentate nucleus and its benefits over atlas-based methods. The Journal of Medical Investigation: JMI, 6(4), 044007.

Nowacki, A., Schlaier, J., Debove, I., & Pollo, C. (2018). Validation of diffusion tensor imaging tractography to visualize the dentatorubrothalamic tract for surgical planning. Journal of Neurosurgery, 130(1), 99–108.

Olchanyi, M. D., Augustinack, J., Haynes, R. L., Lewis, L. D., Cicero, N., Li, J., Destrieux, C., Folkerth, R. D., Kinney, H. C., Fischl, B., Brown, E. N., Iglesias, J. E., & Edlow, B. L. (2025). Automated MRI segmentation of brainstem nuclei critical to consciousness. Human Brain Mapping, 46(14), e70357.

Palay, S. L., & Chan-Palay, V. (2012). Cerebellar Cortex: Cytology and Organization. Springer Science & Business Media.

Palesi, F., De Rinaldis, A., Castellazzi, G., Calamante, F., Muhlert, N., Chard, D., Tournier, J. D., Magenes, G., D’Angelo, E., & Gandini Wheeler-Kingshott, C. A. M. (2017). Contralateral cortico-ponto-cerebellar pathways reconstruction in humans in vivo: implications for reciprocal cerebro-cerebellar structural connectivity in motor and non-motor areas. Scientific Reports, 7(1), 12841.

Palesi, F., Lorenzi, R. M., Casellato, C., Ritter, P., Jirsa, V., Gandini Wheeler-Kingshott, C. A. M., & D’Angelo, E. (2020). The Importance of Cerebellar Connectivity on Simulated Brain Dynamics. Frontiers in Cellular Neuroscience, 14, 240.

Palesi, F., Tournier, J.-D., Calamante, F., Muhlert, N., Castellazzi, G., Chard, D., D’Angelo, E., & Wheeler-Kingshott, C. A. M. (2015). Contralateral cerebello-thalamo-cortical pathways with prominent involvement of associative areas in humans in vivo. Brain Structure & Function, 220(6), 3369–3384.

Petersen, K. J., Reid, J. A., Chakravorti, S., Juttukonda, M. R., Franco, G., Trujillo, P., Stark, A. J., Dawant, B. M., Donahue, M. J., & Claassen, D. O. (2018). Structural and functional connectivity of the nondecussating dentato-rubro-thalamic tract. NeuroImage, 176, 364–371.

Phillips, J. R., Hewedi, D. H., Eissa, A. M., & Moustafa, A. A. (2015). The cerebellum and psychiatric disorders. Frontiers in Public Health, 3, 66.

Radwan, A. M., Sunaert, S., Schilling, K., Descoteaux, M., Landman, B. A., Vandenbulcke, M., Theys, T., Dupont, P., & Emsell, L. (2022). An atlas of white matter anatomy, its variability, and reproducibility based on constrained spherical deconvolution of diffusion MRI. NeuroImage, 254(119029), 119029.

Ramos-Llordén, G., Ning, L., Liao, C., Mukhometzianov, R., Michailovich, O., Setsompop, K., & Rathi, Y. (2020). High-fidelity, accelerated whole-brain submillimeter in vivo diffusion MRI using gSlider-spherical ridgelets (gSlider-SR). Magnetic Resonance in Medicine, 84(4), 1781–1795.

Reddy, C. P., & Rathi, Y. (2016). Joint Multi-Fiber NODDI Parameter Estimation and Tractography Using the Unscented Information Filter. Frontiers in Neuroscience, 10, 166.

Reeber, S. L., Otis, T. S., & Sillitoe, R. V. (2013). New roles for the cerebellum in health and disease. Frontiers in Systems Neuroscience, 7, 83.

Rockland, K. S. (2013). Collateral branching of long-distance cortical projections in monkey: Collateral Branching of Cortical Projections. The Journal of Comparative Neurology, 521(18), 4112–4123.

Rockland, K. S. (2018). Axon collaterals and brain states. Frontiers in Systems Neuroscience, 12, 32.

Rockland, K. S., & Rushmore, R. J. (2025). Cortical white matter: no longer a silent partner. Frontiers in Neuroanatomy, 19(1726067), 1726067.

Rousseau, P.-N., Chakravarty, M. M., & Steele, C. J. (2022). Mapping pontocerebellar connectivity with diffusion MRI. NeuroImage, 264(119684), 119684.

Rushmore, R. J., Bouix, S., Kubicki, M., Rathi, Y., Yeterian, E. H., & Makris, N. (2020). How human is human connectional neuroanatomy? Frontiers in Neuroanatomy, 14, 18.

Rushmore, R. J., Wilson-Braun, P., Papadimitriou, G., Ng, I., Rathi, Y., Zhang, F., O’Donnell, L. J., Kubicki, M., Bouix, S., Yeterian, E., Lemaire, J. J., Calabrese, E., Johnson, G. A., Kikinis, R., & Makris, N. (2020). 3D exploration of the brainstem in 50-micron resolution MRI. Front. Neuroanat. 14, 40. 10.3389/fnana.2020.00040

Sajonz, B. E. A., Frommer, M. L., Reisert, M., Blazhenets, G., Schröter, N., Rau, A., Prokop, T., Reinacher, P. C., Rijntjes, M., Urbach, H., Meyer, P. T., & Coenen, V. A. (2024). Disbalanced recruitment of crossed and uncrossed cerebello-thalamic pathways during deep brain stimulation is predictive of delayed therapy escape in essential tremor. NeuroImage. Clinical, 41(103576), 103576.

Sakai, S. T., Inase, M., & Tanji, J. (1996). Comparison of cerebellothalamic and pallidothalamic projections in the monkey (Macaca fuscata): A double anterograde labeling study. The Journal of Comparative Neurology, 368(2), 215–228.

Sathyanesan, A., Zhou, J., Scafidi, J., Heck, D. H., Sillitoe, R. V., & Gallo, V. (2019). Emerging connections between cerebellar development, behaviour and complex brain disorders. Nature Reviews. Neuroscience, 20(5), 298–313.

Sato, H., & Noda, H. (1991). Divergent axon collaterals from fastigial oculomotor region to mesodiencephalic junction and paramedian pontine reticular formation in macaques. Neuroscience Research, 11(1), 41–54.

Schilling, K. G., Nath, V., Hansen, C., Parvathaneni, P., Blaber, J., Gao, Y., Neher, P., Aydogan, D. B., Shi, Y., Ocampo-Pineda, M., Schiavi, S., Daducci, A., Girard, G., Barakovic, M., Rafael-Patino, J., Romascano, D., Rensonnet, G., Pizzolato, M., Bates, A., … Landman, B. A. (2019). Limits to anatomical accuracy of diffusion tractography using modern approaches. NeuroImage, 185, 1–11.

Schilling, K. G., Petit, L., Rheault, F., Remedios, S., Pierpaoli, C., Anderson, A. W., Landman, B. A., & Descoteaux, M. (2020). Brain connections derived from diffusion MRI tractography can be highly anatomically accurate-if we know where white matter pathways start, where they end, and where they do not go. Brain Structure & Function, 225(8), 2387–2402.

Schilling, K. G., Tax, C. M. W., Rheault, F., Landman, B. A., Anderson, A. W., Descoteaux, M., & Petit, L. (2022). Prevalence of white matter pathways coming into a single white matter voxel orientation: The bottleneck issue in tractography. Human Brain Mapping, 43(4), 1196–1213.

Sereno, M. I., Diedrichsen, J., Tachrount, M., Testa-Silva, G., d’Arceuil, H., & De Zeeuw, C. (2020). The human cerebellum has almost 80% of the surface area of the neocortex. Proceedings of the National Academy of Sciences of the United States of America, 117(32), 19538–19543.

Setsompop, K., Fan, Q., Stockmann, J., Bilgic, B., Huang, S., Cauley, S. F., Nummenmaa, A., Wang, F., Rathi, Y., Witzel, T., & Wald, L. L. (2018). High-resolution in vivo diffusion imaging of the human brain with generalized slice dithered enhanced resolution: Simultaneous multislice (gSlider-SMS). Magnetic Resonance in Medicine, 79(1), 141–151.

Smith, M. C. (1957). The anatomy of the spinocerebellar fibers in man. I. The course of the fibers in the spinal cord and brain stem. The Journal of Comparative Neurology, 108(2), 285–352.

Smith, R. E., Tournier, J.-D., Calamante, F., & Connelly, A. (2012). Anatomically-constrained tractography: improved diffusion MRI streamlines tractography through effective use of anatomical information. NeuroImage, 62(3), 1924–1938.

Somana, R., & Walberg, F. (1978). Cerebellar afferents from the paramedian reticular nucleus studied with retrograde transport of horseradish peroxidase. Anatomy and Embryology, 154(3), 353–368.

Steele, C. J., Anwander, A., Bazin, P.-L., Trampel, R., Schaefer, A., Turner, R., Ramnani, N., & Villringer, A. (2017). Human Cerebellar Sub-millimeter Diffusion Imaging Reveals the Motor and Non-motor Topography of the Dentate Nucleus. Cerebral Cortex, 27(9), 4537–4548.

Strick, P. L., Dum, R. P., & Fiez, J. A. (2009). Cerebellum and nonmotor function. Annual Review of Neuroscience, 32, 413–434.

Sugimoto, T., Mizuno, N., & Itoh, K. (1981). An autoradiographic study on the terminal distribution of cerebellothalamic fibers in the cat. Brain Research, 215(1-2), 29–47.

Takahashi, E., Song, J. W., Folkerth, R. D., Grant, P. E., & Schmahmann, J. D. (2013). Detection of postmortem human cerebellar cortex and white matter pathways using high angular resolution diffusion tractography: a feasibility study. NeuroImage, 68, 105–111.

Tang, Y., Sun, W., Toga, A. W., Ringman, J. M., & Shi, Y. (2018). A probabilistic atlas of human brainstem pathways based on connectome imaging data. NeuroImage, 169, 227–239.

Tchetchenian, A., Zekelman, L., Chen, Y., Rushmore, J., Zhang, F., Yeterian, E. H., Makris, N., Rathi, Y., Meijering, E., Song, Y., & O’Donnell, L. J. (2024). Deep multimodal saliency parcellation of cerebellar pathways: Linking microstructure and individual function through explainable multitask learning. Human Brain Mapping, 45(12), e70008.

ten Donkelaar, H. J., den Dunnen, W., van de Warrenburg, B., Lammens, M., & Wesseling, P. (2020). The cerebellum. In Clinical Neuroanatomy (pp. 539–589). Springer International Publishing.

Tolbert, D. L., & Bantli, H. (1979). An HRP and autoradiographic study of cerebellar corticonuclear-nucleocortical reciprocity in the monkey. Experimental Brain Research, 36(3), 563–571.

Tolbert, D. L., Bantli, H., & Bloedel, J. R. (1978). Organizational features of the cat and monkey cerebellar nucleocortical projection. The Journal of Comparative Neurology, 182(1), 39–56.

Tournier, J.-D., Smith, R., Raffelt, D., Tabbara, R., Dhollander, T., Pietsch, M., Christiaens, D., Jeurissen, B., Yeh, C.-H., & Connelly, A. (2019). MRtrix3: A fast, flexible and open software framework for medical image processing and visualisation. NeuroImage, 202(116137), 116137.

van Baarsen, K. M., Kleinnijenhuis, M., Jbabdi, S., Sotiropoulos, S. N., Grotenhuis, J. A., & van Cappellen van Walsum, A. M. (2016). A probabilistic atlas of the cerebellar white matter. NeuroImage, 124(Pt A), 724–732.

Voogd, J. (2003). The human cerebellum. Journal of Chemical Neuroanatomy, 26(4), 243–252.

Voogd, J., Gerrits, N. M., & Hess, D. T. (1987). Parasagittal zonation of the cerebellum in macaques: An analysis based on acetylcholinesterase histochemistry. In Cerebellum and Neuronal Plasticity (pp. 15–40). Springer US.

Wang, B., LeBel, A., & D’Mello, A. M. (2025). Ignoring the cerebellum is hindering progress in neuroscience. Trends in Cognitive Sciences, 29(4), 318–330.

Wang, X., Wang, J., Chen, Y., O’Donnell, L. J., & Zhang, F. (2026). Bridging modalities: Joint synthesis and registration framework for aligning diffusion MRI with T1-weighted images. In arXiv [eess.IV]. arXiv. 10.48550/arXiv.2601.11689

Warrington, S., Thompson, E., Bastiani, M., Dubois, J., Baxter, L., Slater, R., Jbabdi, S., Mars, R. B., & Sotiropoulos, S. N. (2022). Concurrent mapping of brain ontogeny and phylogeny within a common space: Standardized tractography and applications. Science Advances, 8(42), eabq2022.

Wassermann, D., Makris, N., Rathi, Y., Shenton, M., Kikinis, R., Kubicki, M., & Westin, C.-F. (2016). The white matter query language: a novel approach for describing human white matter anatomy. Brain Structure & Function, 221(9), 4705–4721.

Wu, T., & Hallett, M. (2013). The cerebellum in Parkinson’s disease. Brain: A Journal of Neurology, 136(Pt 3), 696–709.

Wyatt, K. D., Tanapat, P., & Wang, S. S.-H. (2005). Speed limits in the cerebellum: constraints from myelinated and unmyelinated parallel fibers. The European Journal of Neuroscience, 21(8), 2285–2290.

Xu, X., Chen, Y., Zekelman, L., Rathi, Y., Makris, N., Zhang, F., & O’Donnell, L. J. (2023). Tractography-based parcellation of cerebellar dentate nuclei via a deep nonnegative matrix factorization clustering method. 2023 IEEE 20th International Symposium on Biomedical Imaging *(ISBI)*, 1–4.

Yamada, J., & Noda, H. (1987). Afferent and efferent connections of the oculomotor cerebellar vermis in the macaque monkey. In The Journal of Comparative Neurology (Vol. 265, Issue 2, pp. 224–241). 10.1002/cne.902650207

Yeh, C.-H., Jones, D. K., Liang, X., Descoteaux, M., & Connelly, A. (2020). Mapping Structural Connectivity Using Diffusion MRI: Challenges and Opportunities. Journal of Magnetic Resonance Imaging: JMRI. 10.1002/jmri.27188

Yeh, F.-C., Panesar, S., Fernandes, D., Meola, A., Yoshino, M., Fernandez-Miranda, J. C., Vettel, J. M., & Verstynen, T. (2018). Population-averaged atlas of the macroscale human structural connectome and its network topology. NeuroImage, 178, 57–68.

Yin, H., Zong, F., Deng, X., Zhang, D., Zhang, Y., Wang, S., Wang, Y., & Zhao, J. (2023). The language-related cerebro-cerebellar pathway in humans: a diffusion imaging-based tractographic study. Quantitative Imaging in Medicine and Surgery, 13(3), 1399–1416.

Zekelman, L. R., Cetin-Karayumak, S., Chen, Y., Almeida, M., Legarreta, J. H., Rushmore, J., Pieper, S., Lan, Z., Desmond, J. E., Baird, L. C., Makris, N., Rathi, Y., Zhang, F., Golby, A. J., & O’Donnell, L. J. (2025). Consistent cerebellar pathway-cognition associations across pre-adolescents & young adults: a diffusion MRI study of 9000+ participants. In bioRxivorg. 10.1101/2025.02.05.636737

Zhang, F., Breger, A., Cho, K. I. K., Ning, L., Westin, C.-F., O’Donnell, L. J., & Pasternak, O. (2021). Deep learning based segmentation of brain tissue from diffusion MRI. NeuroImage, 233, 117934.

Zhang, F., Chen, Y., Ning, L., Rushmore, J., Liu, Q., Du, M., Hassanzadeh-Behbahani, S., Legarreta, J. H., Yeterian, E., Makris, N., Rathi, Y., & O’Donnell, L. J. (2024). Assessment of the depiction of superficial white matter using ultra-high-resolution diffusion MRI. Human Brain Mapping, 45(14), e70041.

Zhang, F., Théberge, A., Jodoin, P.-M., Descoteaux, M., & O’Donnell, L. J. (2025). Think deep in the tractography game: deep learning for tractography computing and analysis. Brain Structure & Function, 230(6), 100.

Zhang, F., Wu, Y., Norton, I., Rigolo, L., Rathi, Y., Makris, N., & O’Donnell, L. J. (2018). An anatomically curated fiber clustering white matter atlas for consistent white matter tract parcellation across the lifespan. NeuroImage, 179, 429–447.

Zhang, Z., Descoteaux, M., Zhang, J., Girard, G., Chamberland, M., Dunson, D., Srivastava, A., & Zhu, H. (2018). Mapping population-based structural connectomes. NeuroImage, 172, 130–145.

Zheng, J., Yang, Q., Makris, N., Huang, K., Liang, J., Ye, C., Yu, X., Tian, M., Ma, T., Mou, T., Guo, W., Kikinis, R., & Gao, Y. (2023). Three-dimensional digital reconstruction of the cerebellar cortex: Lobule thickness, surface area measurements, and layer architecture. *Cerebellum (London*, England), 22(2), 249–260.

