## Supplementary materials for "Anatomically constrained and curated cerebellar tractography (ACCURAT): an open framework and a pathway-specific neuroanatomical reference"

[1] Department of Radiology, Brigham and Women's Hospital, Mass General Brigham, Boston MA, USA [2] Harvard Medical School, Boston MA, USA [3] Department of Anatomy and Neurobiology, Boston University Chobanian & Avedisian School of Medicine, Boston MA, USA [4] Department of Psychiatry, Brigham and Women's Hospital, Mass General Brigham, Boston MA, USA [5] Department of Psychology, Colby College, Waterville ME, USA

### Supplementary Materials

#### Expert segmentation of nuclei and other ROIs

Multiple ROIs were successfully segmented to enable the ACCURAT framework. We provide volumetric information for each in Table S1, including, for reference, the voxel count at the equivalent resolution of the Human Connectome Project Young Adult (HCP-YA) dataset (1.25 mm isotropic).

**Table S1.** Volumetric characteristics of anatomical regions used for ACCURAT (mean ± SD).

| ROI | Category | Volume (mm³, mean ± SD) | # Voxels (0.76 mm iso) | # Voxels (1.25 mm iso equivalent) |
|---|---|---|---|---|
| **Dentate nucleus** | Deep cerebellar nuclei | 1603.6 ± 349.53 | 3666 ± 799 | 821 ± 178 |
| **Emboliform nucleus** | Deep cerebellar nuclei | 89.71 ± 16.73 | 205 ± 38 | 46 ± 9 |
| **Globose nucleus** | Deep cerebellar nuclei | 28.28 ± 19.16 | 65 ± 44 | 15 ± 10 |
| **Fastigial nucleus** | Deep cerebellar nuclei | 85.39 ± 16.24 | 195 ± 37 | 43 ± 8 |
| **Inferior olivary nucleus** | Brainstem nuclei | 616.81 ± 98.65 | 1410 ± 226 | 316 ± 51 |
| **Red nucleus** | Brainstem nuclei | 471.01 ± 78.97 | 1077 ± 181 | 241 ± 40 |
| **Basilar pons** | Brainstem region | 8010.93 ± 1116.49 | 18316 ± 2553 | 4101 ± 572 |
| **Pontine tegmentum** | Brainstem region | 3289.66 ± 471.57 | 7521 ± 1078 | 1684 ± 241 |
| **ICP ROI** | Brainstem region | 278.27 ± 54.06 | 636 ± 124 | 142 ± 28 |

#### Quantitative reconstruction with stricter threshold

Table S2 shows the number of subjects in which each bundle was successfully identified with at least 10 streamlines for both PTT and UKF data. Results are consistent across pathways and tractography methods, with the olivocerebellar ICP pathway showing a reduced extraction rate on PTT data, due to the increased difficulty of reconstruction.

**Table S2**. Number of subjects with successful bundle reconstruction (≥5 or ≥10 streamlines; L: left; R: right; B: both hemispheres).

| | PTT | | UKF | |
|---|---|---|---|---|
| **Pathway** | **# Subjects with ≥5 streamlines** | **# Subjects with ≥10 streamlines** | **# Subjects with ≥5 streamlines** | **# Subjects with ≥10 streamlines** |
| | 9/9 | 9/9 | 9/9 | 9/9 |
| ***Extrinsic cerebellar pathways*** | | | | |
| **ICP olivocerebellar** | L: 6; R: 7; B: 6 | L: 3; R: 6; B: 3 | L: 9; R: 8; B: 8 | L: 9; R: 8; B: 8 |
| **ICP peduncle** | L: 9; R: 9; B: 9 | L: 9; R: 9; B: 9 | L: 9; R: 9; B: 9 | L: 9; R: 9; B: 9 |
| **MCP pontocerebellar** | L: 9; R: 9; B: 9 | L: 9; R: 9; B: 9 | L: 9; R: 9; B: 9 | L: 9; R: 9; B: 9 |
| **MCP reticulocerebellar** | L: 9; R: 9; B: 9 | L: 9; R: 9; B: 9 | L: 9; R: 9; B: 9 | L: 9; R: 9; B: 9 |
| **SCP dentato-rubral** | L: 9; R: 9; B: 9 | L: 9; R: 9; B: 9 | L: 9; R: 9; B: 9 | L: 9; R: 9; B: 9 |
| **SCP dentato-rubro-thalamic** | L: 9; R: 9; B: 9 | L: 9; R: 9; B: 9 | L: 9; R: 9; B: 9 | L: 9; R: 9; B: 9 |
| **SCP dentato-thalamic** | L: 9; R: 9; B: 9 | L: 9; R: 9; B: 9 | L: 9; R: 9; B: 9 | L: 9; R: 9; B: 9 |
| **SCP dentato-olivary** | L: 9; R: 9; B: 9 | L: 9; R: 9; B: 9 | L: 9; R: 9; B: 9 | L: 9; R: 8; B: 8 |
| ***Intrinsic cerebellar pathways*** | | | | |
| **Purkinje cortico-dentate** | L: 9; R: 9; B: 9 | L: 9; R: 9; B: 9 | L: 9; R: 9; B: 9 | L: 9; R: 9; B: 9 |
| **Purkinje cortico-emboliform** | L: 9; R: 9; B: 9 | L: 9; R: 9; B: 9 | L: 9; R: 9; B: 9 | L: 9; R: 9; B: 9 |
| **Purkinje cortico-globose** | L: 9; R: 9; B: 9 | L: 9; R: 9; B: 9 | L: 9; R: 9; B: 9 | L: 9; R: 9; B: 9 |
| **Purkinje cortico-fastigial** | L: 9; R: 9; B: 9 | L: 9; R: 9; B: 9 | L: 9; R: 9; B: 9 | L: 9; R: 9; B: 9 |
| **Parallel** | L: 9; R: 9; B: 9 | L: 9; R: 9; B: 9 | L: 9; R: 9; B: 9 | L: 9; R: 9; B: 9 |

#### Additional visual results of ACCURAT bundle extraction

Figures S1-S5 show the bundles of interest extracted from UKF tractography data on the same participant shown across Figures 4-9.

**UKF pathways for the same participant shown previously**

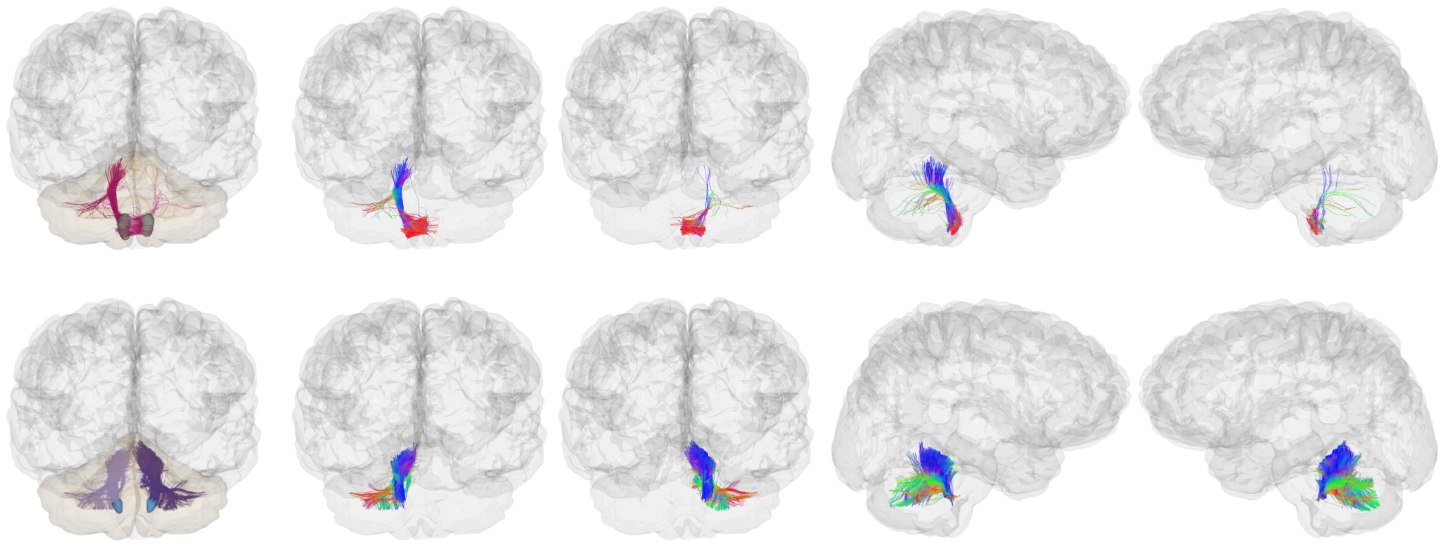

**Figure S1. Inferior cerebellar peduncles as extracted by ACCURAT for the same exemplar participant shown previously, using UKF data**. Top row: olivocerebellar pathway; bottom row: inferior cerebellar peduncle.

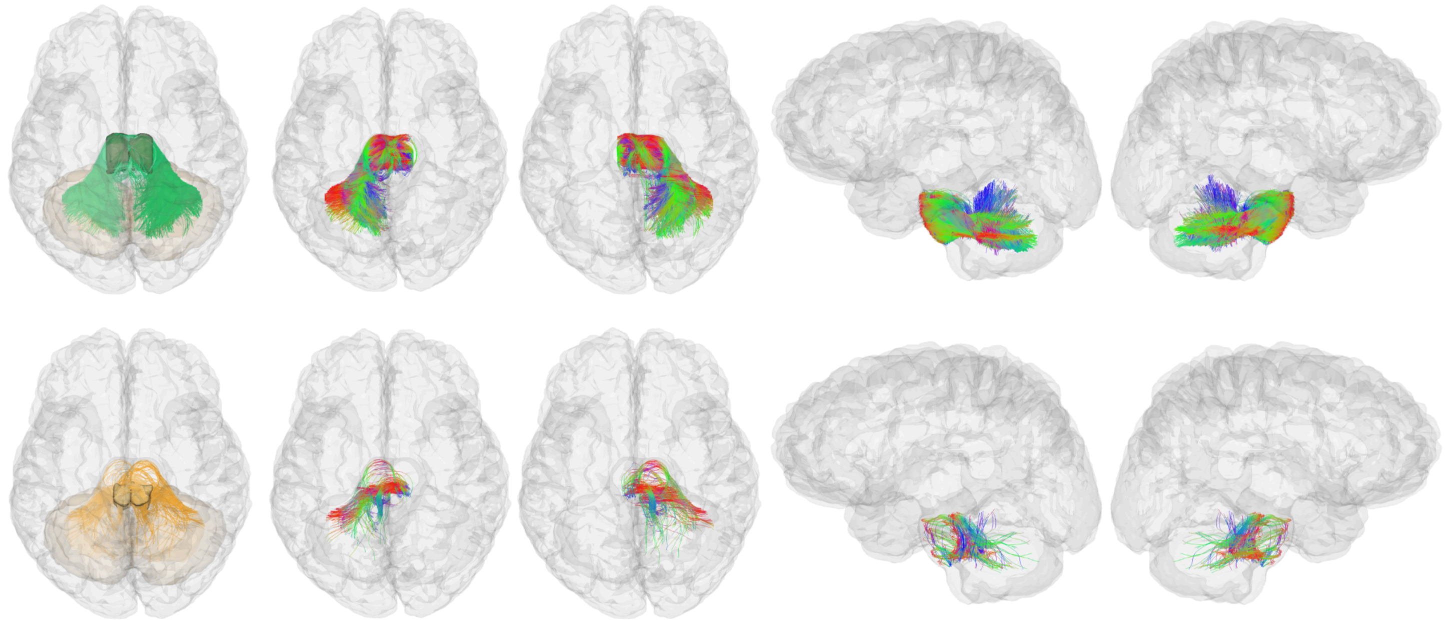

**Figure S2. Middle cerebellar peduncles as extracted by ACCURAT for the same exemplar participant shown previously, using UKF data.** Top row: pontocerebellar; bottom row: pontine reticulocerebellar.

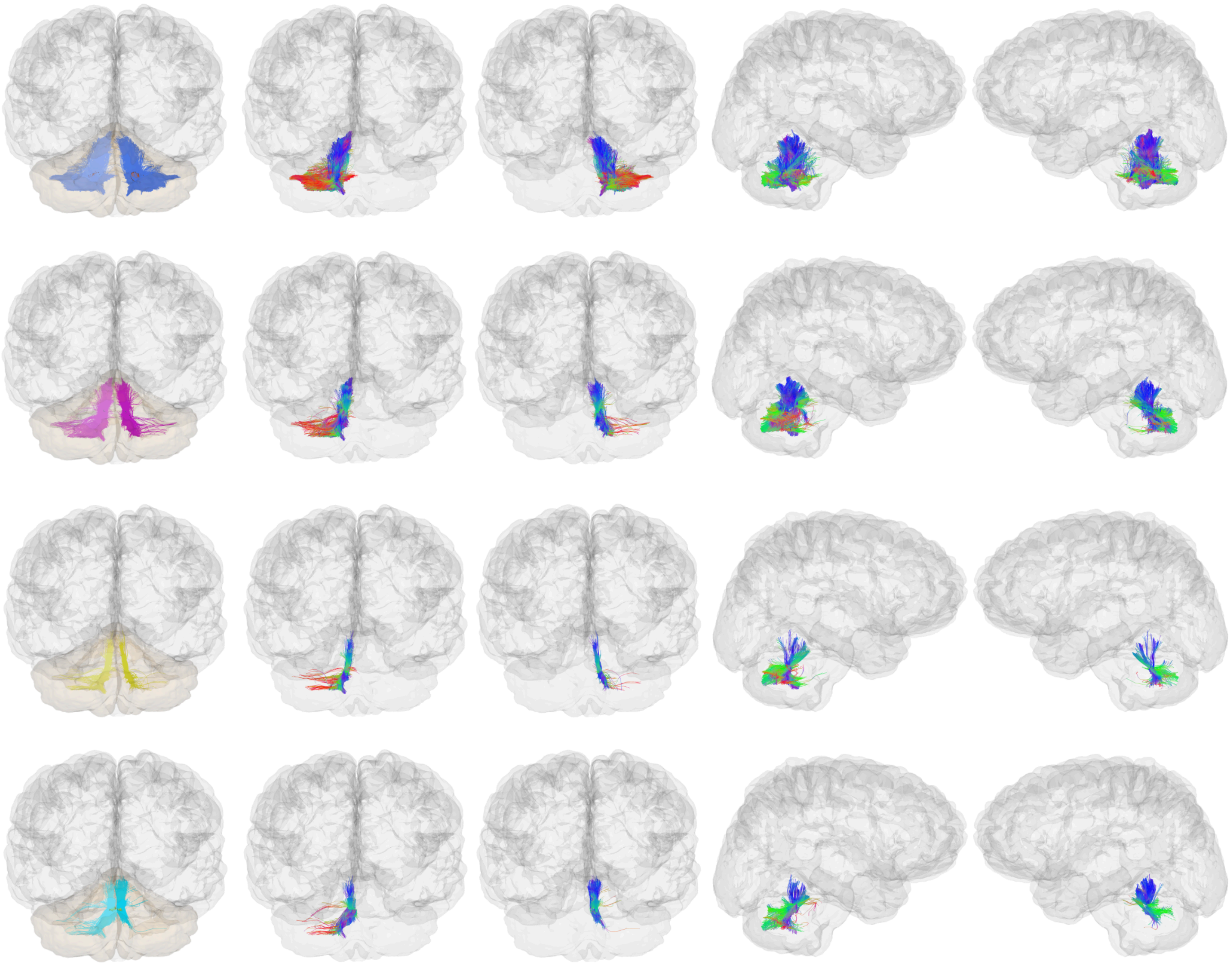

**Figure S3. Purkinje corticonuclear projections as extracted by ACCURAT for the same exemplar participant shown previously, using UKF data**. Top row: Purkinje cortico-dentate; second row: Purkinje cortico-emboliform; third row: Purkinje cortico-globose; bottom row: Purkinje cortico-fastigial.

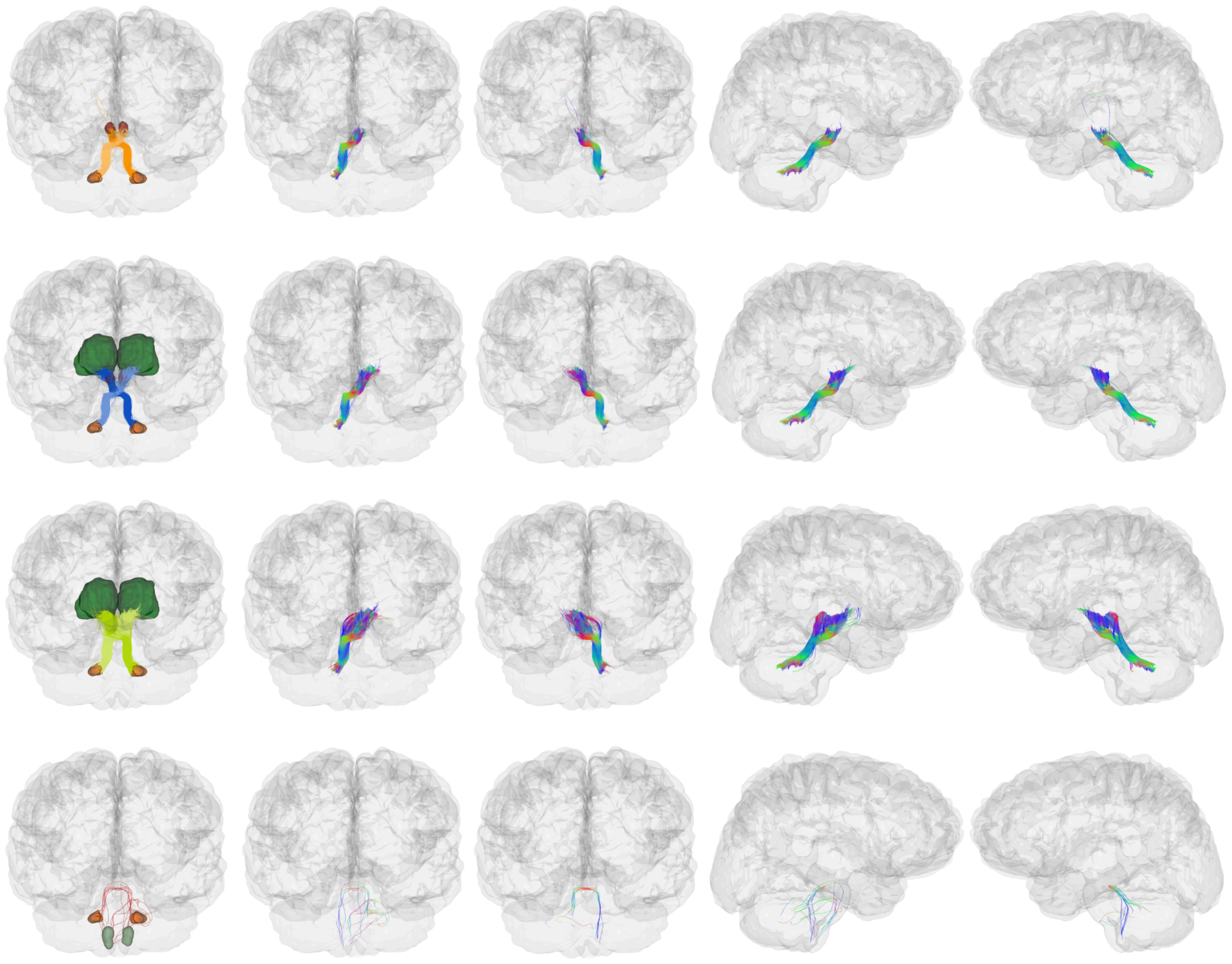

**Figure S4. Superior cerebellar peduncles as extracted by ACCURAT for the same exemplar participant shown previously, using UKF data.** Top row: dentato-rubral; second row: dentato-rubro-thalamic; third row: dentato-thalamic; bottom row: dentato-olivary.

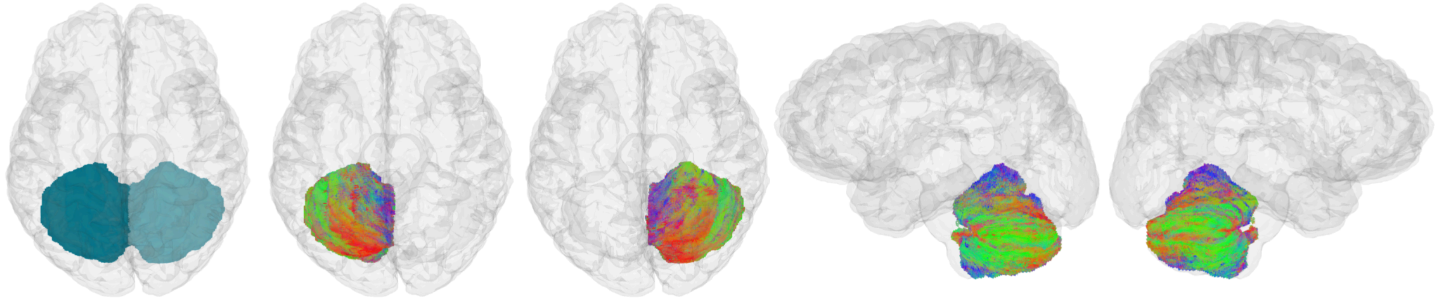

**Figure S5. Parallel fibers as extracted by ACCURAT for the same exemplar participant shown previously, using UKF data.**

**Pathways for an additional participant**

Figures S6-S15 show the pathways of interest for an additional participant extracted from PTT (Figures S6-S10) and UKF data (Figures S11-S15).

*PTT pathways*

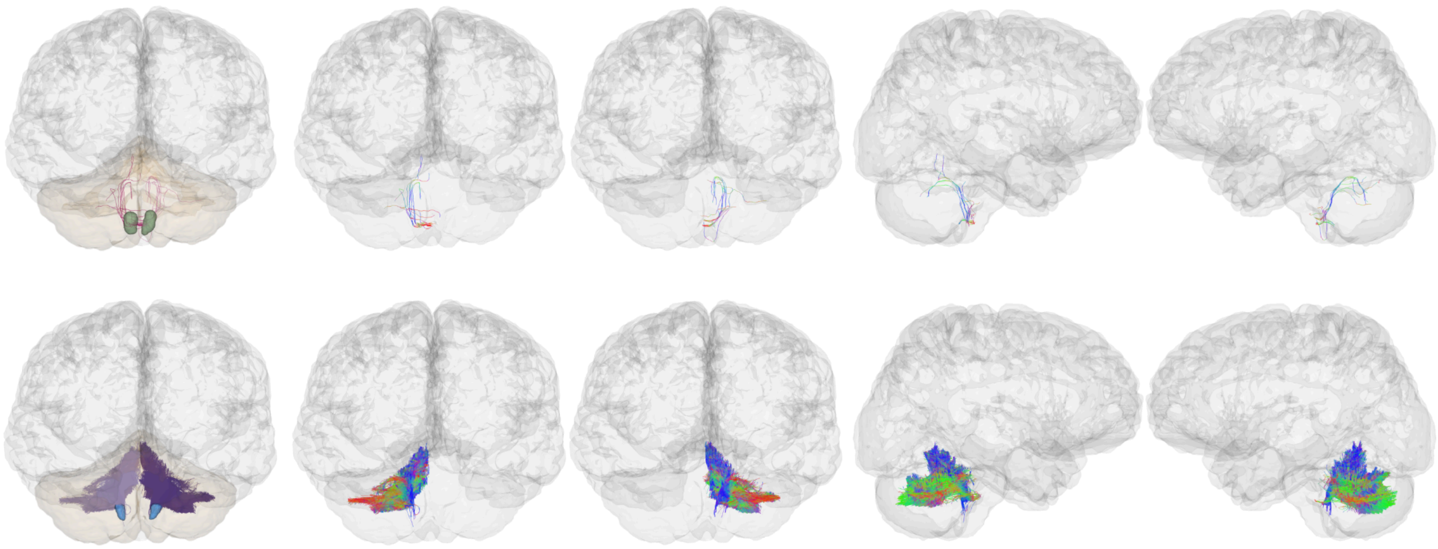

**Figure S6. Inferior cerebellar peduncles as extracted by ACCURAT for an additional exemplar participant, using PTT data.** Top row: olivocerebellar pathway; bottom row: inferior cerebellar peduncle.

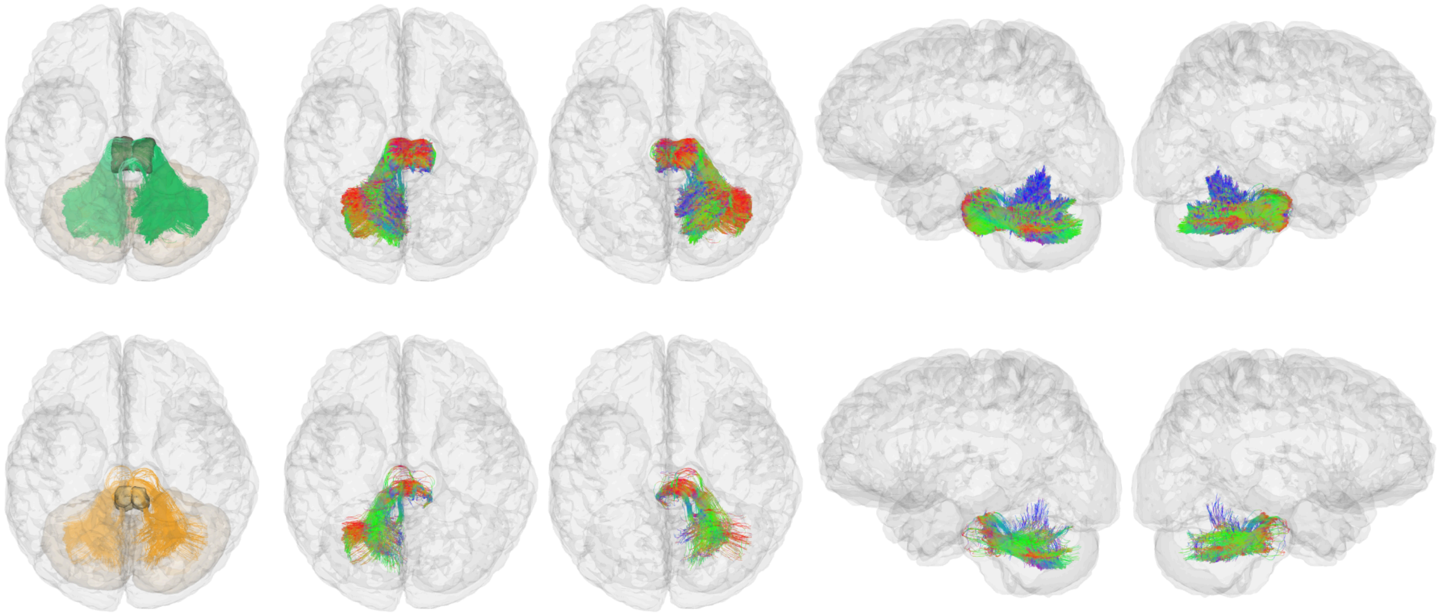

**Figure S7. Middle cerebellar peduncles as extracted by ACCURAT for an additional exemplar participant, using PTT data.** Top row: pontocerebellar; bottom row: pontine reticulocerebellar.

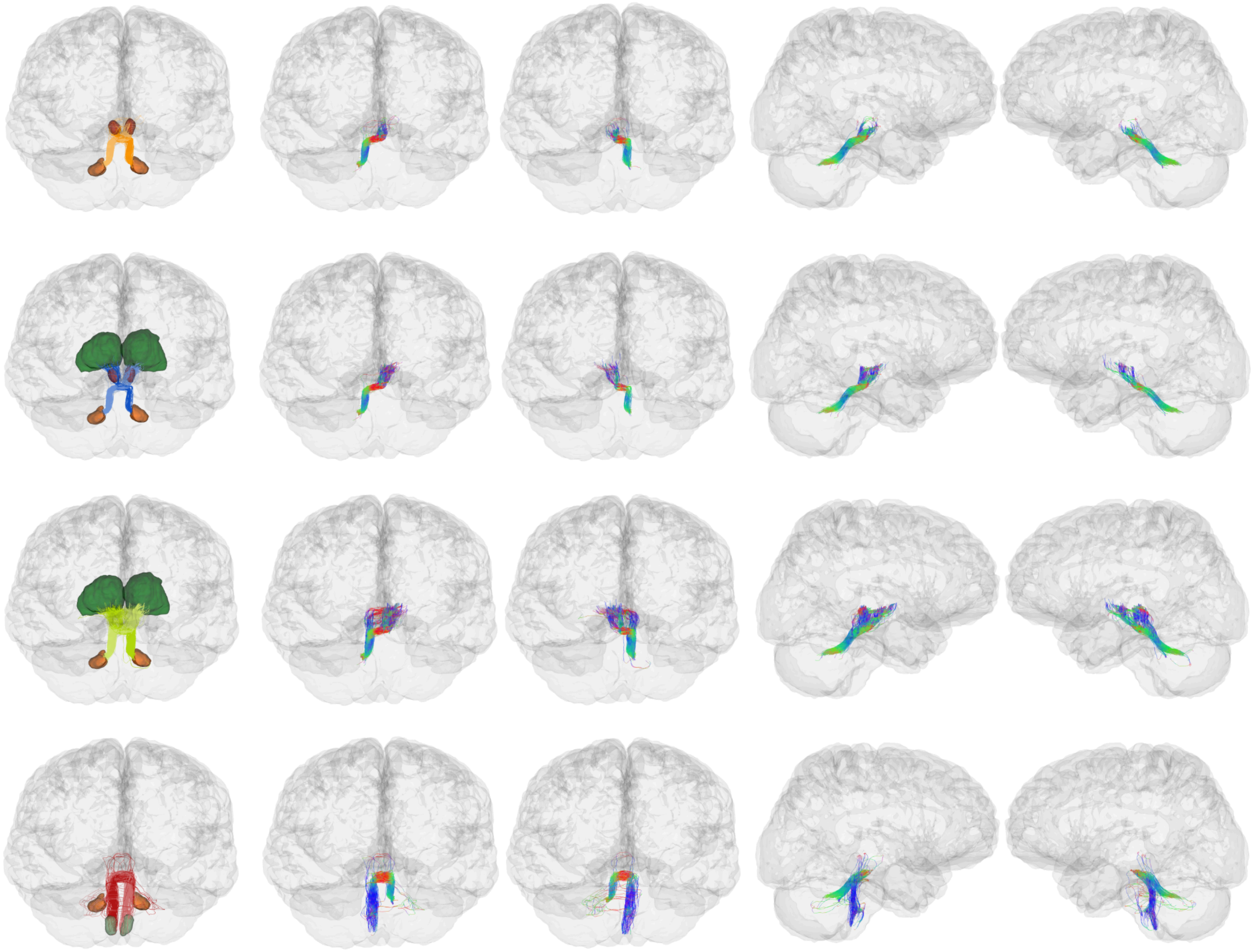

**Figure S8. Superior cerebellar peduncles projections as extracted by ACCURAT for an additional exemplar participant, using PTT data**. Top row: dentato-rubral; second row: dentato-rubro-thalamic; third row: dentato-thalamic; bottom row: dentato-olivary.

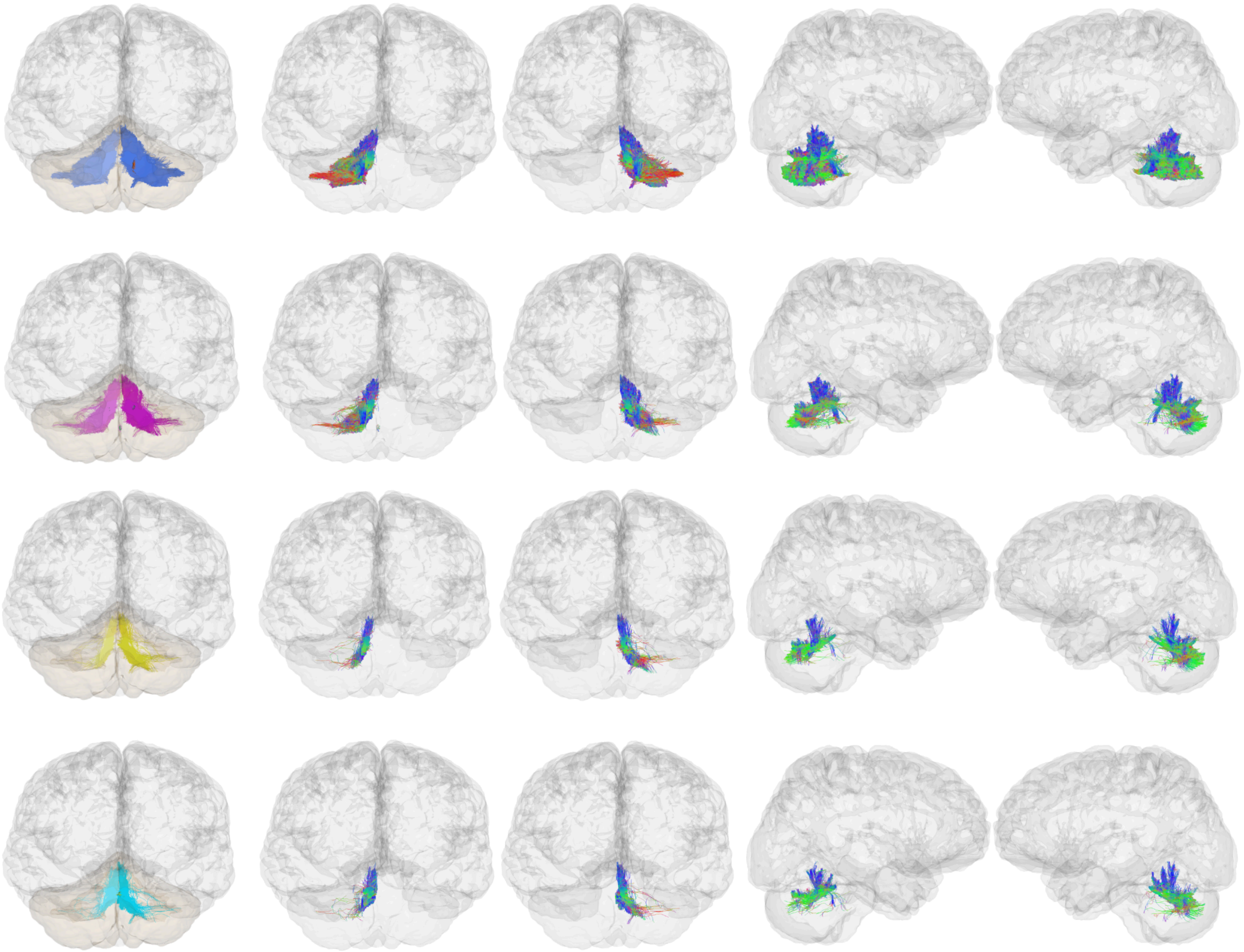

**Figure S9. Purkinje corticonuclear projections as extracted by ACCURAT for an additional exemplar participant, using PTT data.** Top row: Purkinje cortico-dentate; second row: Purkinje cortico-emboliform; third row: Purkinje cortico-globose; bottom row: Purkinje cortico-fastigial.

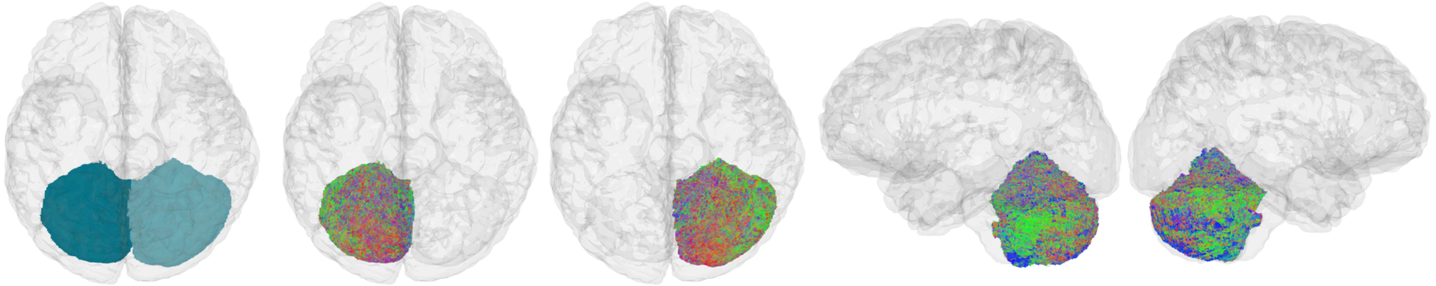

**Figure S10. Parallel fibers as extracted by ACCURAT for an additional exemplar participant, using PTT data.**

*UKF pathways*

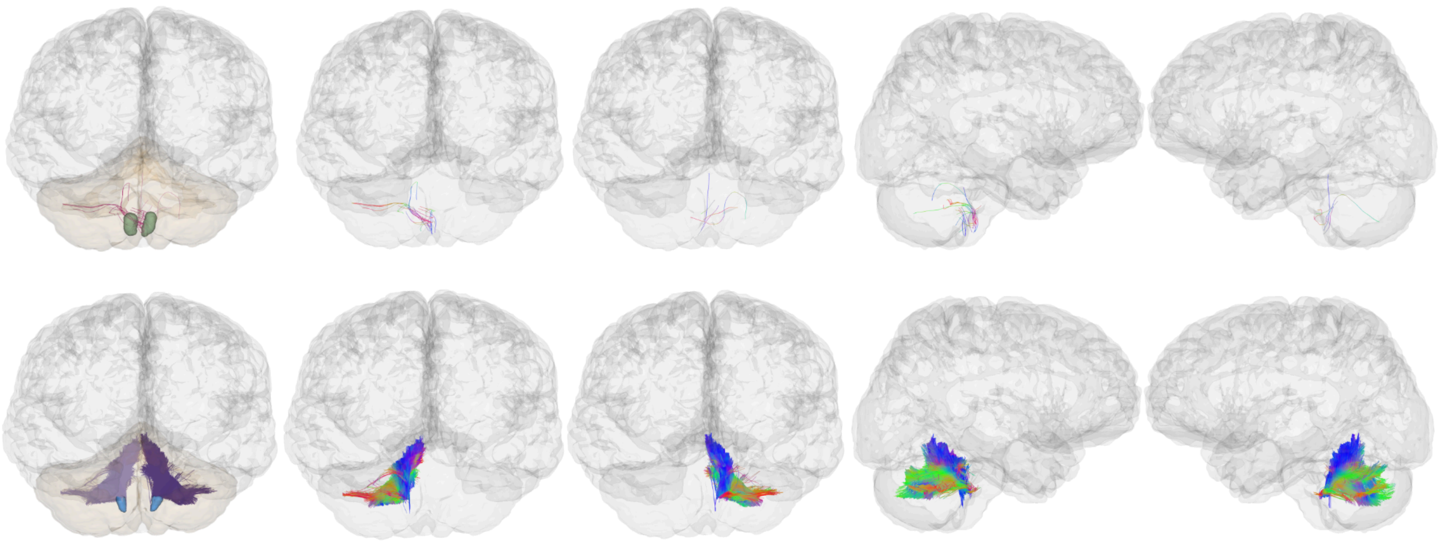

**Figure S11. Inferior cerebellar peduncles as extracted by ACCURAT for an additional exemplar participant, using UKF data.** Top row: olivocerebellar pathway; bottom row: inferior cerebellar peduncle.

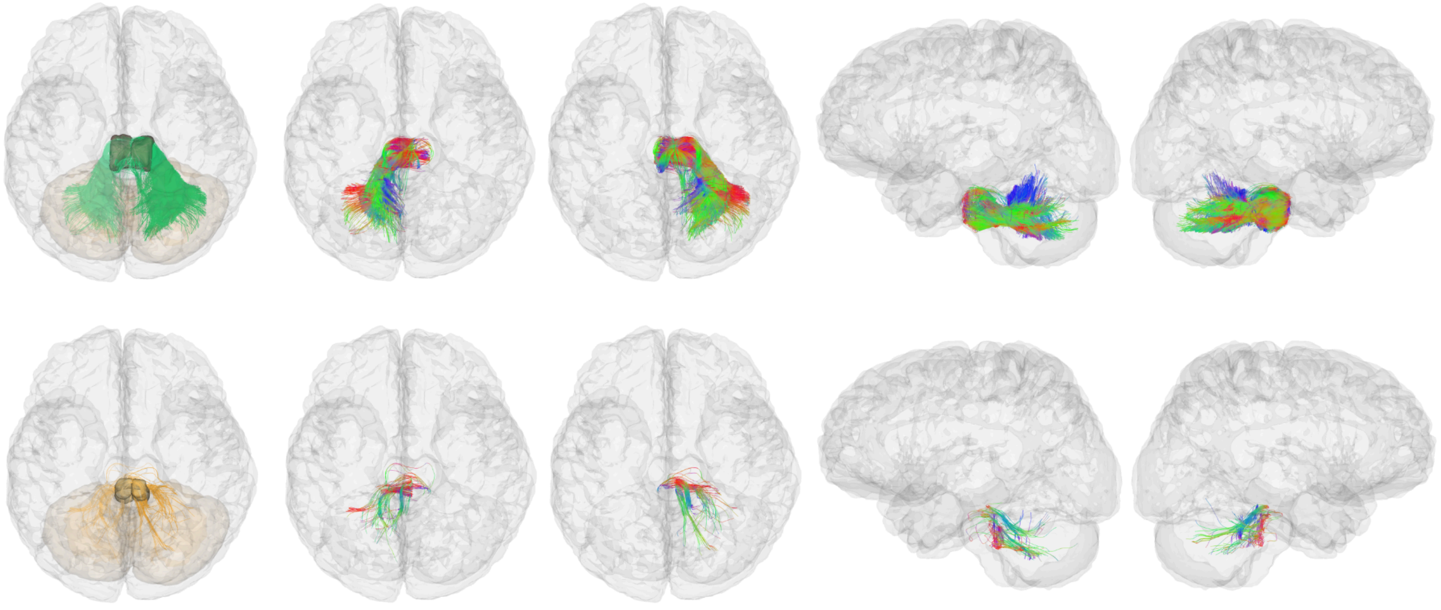

**Figure S12. Middle cerebellar peduncles as extracted by ACCURAT for an additional exemplar participant, using UKF data.** Top row: pontocerebellar; bottom row: pontine reticulocerebellar.

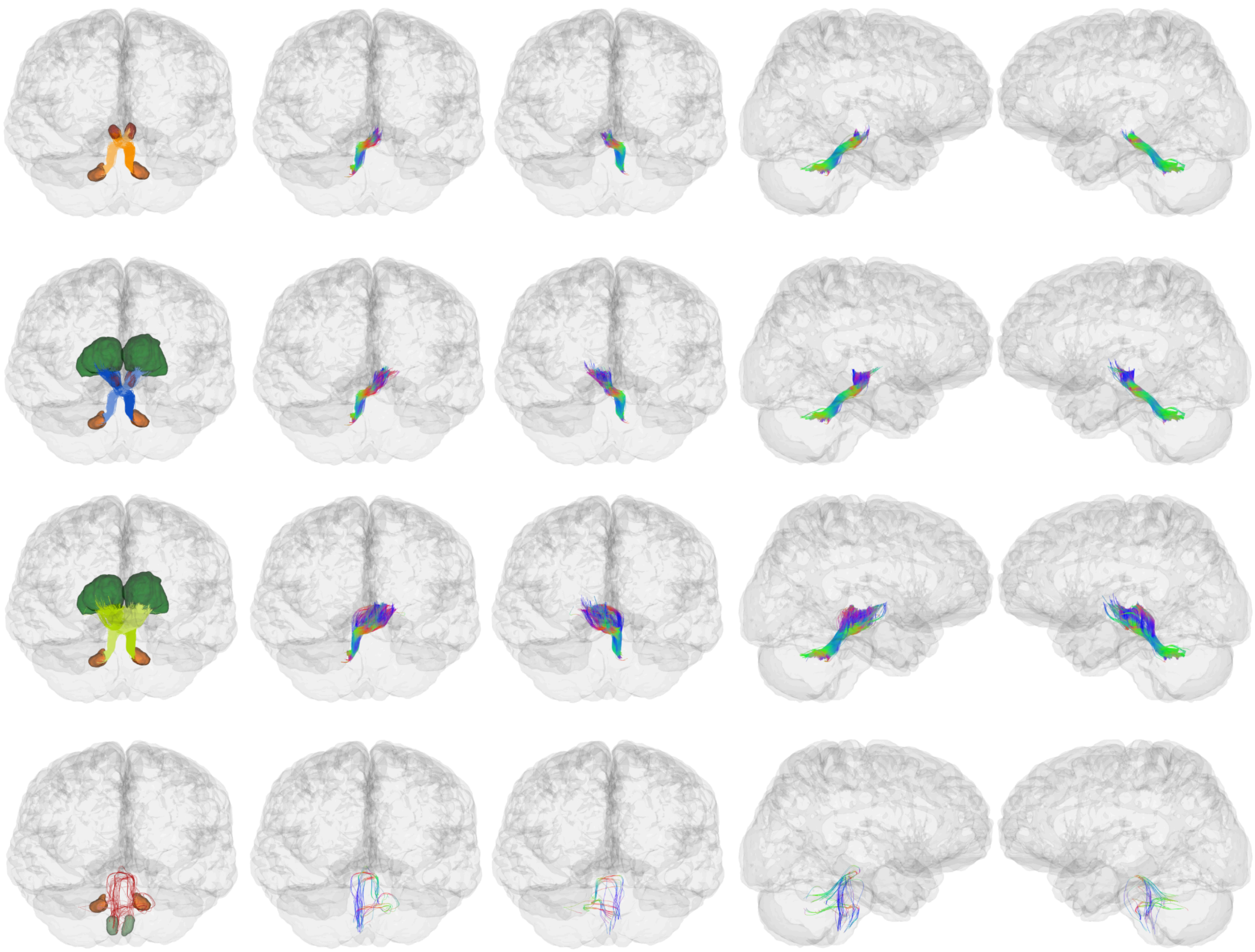

**Figure S13. Superior cerebellar peduncles projections as extracted by ACCURAT for an additional exemplar participant on UKF data**. Top row: dentato-rubral; second row: dentato-rubro-thalamic; third row: dentato-thalamic; bottom row: dentato-olivary.

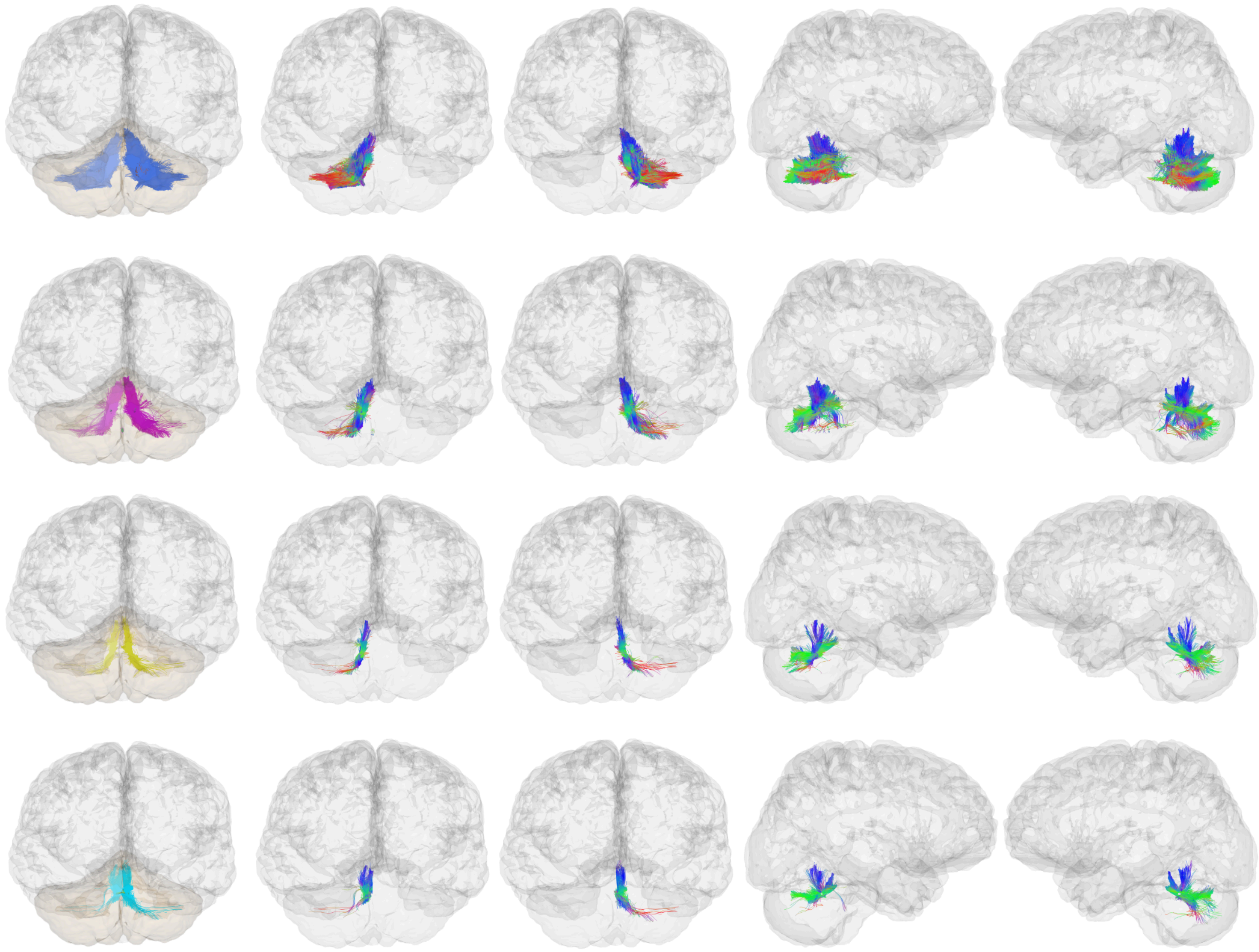

**Figure S14. Purkinje corticonuclear projections as extracted by ACCURAT for an additional exemplar participant on UKF data.** Top row: Purkinje cortico-dentate; second row: Purkinje cortico-emboliform; third row: Purkinje cortico-globose; bottom row: Purkinje cortico-fastigial.

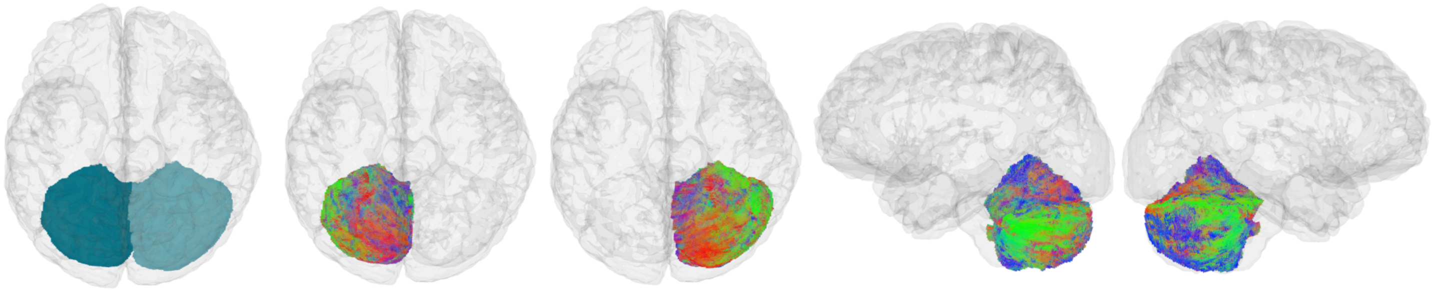

**Figure S15. Parallel fibers as extracted by ACCURAT for an additional exemplar participant on UKF data.**

### Supplementary Methods

#### Supplementary Methods S1 — Diffusion MRI acquisition

Diffusion MRI was acquired using the gSlider sequence (Ramos-Llordén et al., 2020; Setsompop et al., 2018). Acquisition parameters were TE = 87 ms, TR = 3500 ms, in-plane acceleration ×3, multiband acceleration ×2, and 6/8 partial-Fourier encoding. Spatial resolution was 0.76 × 0.76 × 3.8 mm with five RF encodings, which were used to reconstruct isotropic 0.76 mm voxels.

Diffusion gradients were applied along 60 directions at b = 1000 s/mm², with three non-diffusion volumes and a reverse phase-encoding acquisition for EPI distortion correction. The acquisition was repeated three times, resulting in a total scan time of approximately one hour.

The repeated acquisitions were combined using the gSlider reconstruction to produce the final diffusion dataset.

---

#### Supplementary Methods S2 — Preprocessing

Distortions due to susceptibility, eddy currents, and head motion were corrected using FSL's *topup* and *eddy* tools (Jenkinson et al., 2012). Because the acquisition included repeated scans, all acquisitions were merged prior to correction to enable motion and eddy-current estimation across the full dataset.

Spherical harmonic basis functions were fit to each scan, and signals were averaged on a canonical set of gradient directions to obtain the final reconstructed gSlider dataset. This averaging improved signal-to-noise ratio by a factor of √3.

---

### Supplementary Methods S3 — Tractography parameters

Parameters were adjusted empirically to provide reasonable streamline support for the bundles of interest.

##### Parallel Transport Tractography (PTT)

White matter and cerebrospinal fluid response functions were estimated using the Dhollander method and used to compute fiber orientation distributions using multi-shell multi-tissue constrained spherical deconvolution in MRtrix3 (Tournier et al., 2019). Probabilistic tractography was then performed using the parallel transport tractography (PTT) algorithm (Aydogan & Shi, 2021).

Tractography parameters included a minimum streamline length of 6 mm, maximum streamline length of 340 mm, step size of 0.01 mm, probe length of 0.1 mm, probe radius of 0.2 mm, minimum radius of curvature of 0.2 mm, and minimum FOD data support of 0.03.

A total of 3.5 million streamlines were generated.

##### Unscented Kalman Filter (UKF) tractography

Multi-tensor tractography was performed using the unscented Kalman filter (UKF) approach (Reddy & Rathi, 2016) with an FA seeding threshold of 0.1, stopping FA threshold of 0.08, stopping DWI signal threshold of 0.06, three seeds per voxel, and a step size of 0.3 mm.

A total of 2.28 ± 0.2 million streamlines were generated.
